# High-throughput platforms for genetic perturbation screening using CRISPR/Cas9 in human iPSC-derived macrophages for drug discovery

**DOI:** 10.64898/2026.09.11.750978

**Authors:** Matteo Martufi, Eleni Karagianni, Tasos Papanikos, Tim Ashlin, Aikaterini Geladaki, Benedetta Carbone, Katja Remlinger, Andres Quintero Moreno, William Pembroke, Charlie Haslam, Nico Zinn, Denise Vlachou, Margarida Almeida, Dimitrios Tsaousis, Stephan Gade, Bin Sun, Muhammad Maqbool, Holger Franken, Frederik Ziebell, Alessandro Di Tullio, Lewis Entwistle, Naren Srinivasan, Mercedes Lobera, Lisa Mohamet

**Affiliations:** GlaxoSmithKline R&D, Gunnels Wood Rd, Hertfordshire, Stevenage SG1 2NY, UK; GlaxoSmithKline R&D, Meyerhofstrasse 1, 69117 Heidelberg, Germany; GlaxoSmithKline R&D, 1250S Collegeville Rd, Collegeville, PA, 19426 USA; GlaxoSmithKline R&D, 200 Cambridge Park Drive, Cambridge, MA, 02140, USA

## Abstract

Human induced Pluripotent Stem Cell (hiPSC) models have revolutionised drug discovery, offering high translational relevance for recapitulating disease biology and thereby the potential to help reduce drug attrition. Macrophages are pivotal for maintaining tissue homeostasis and orchestrating immune responses; their dysregulation underpins several diseases, including autoinflammatory disorders, neurodegeneration, and cancer. Therapeutically targeting this cell type presents an attractive strategy to simultaneously influence multiple cellular mediators and functions. We have established a scalable, semi-automated, and physiologically relevant hiPSC-derived macrophage model, rigorously characterised through deep comparative multi-omics. We have also integrated our hiPSC-derived macrophage platform with large-scale CRISPR screening workflows designed for parallel genetic interrogation of thousands of gene targets, in both arrayed and pooled formats. In this manuscript, we apply those genetic screening methods to hiPSC-derived macrophages and showcase how genetic perturbations alter pro- and anti-inflammatory transcriptional signatures and significantly impact functional phenotypes in this cell model. This integrated approach allows for the exploration of novel genetic insights linked to causal disease biology, advancing myeloid cell-associated target discovery across a broad spectrum of therapeutic areas.

## Introduction

The innate immune system serves as the body’s primary defence against tissue damage and infection. Macrophages are among the earliest cellular responders, executing diverse functions crucial for restoring tissue homeostasis, including phagocytosis, antigen presentation, and cytokine secretion. Proper macrophage function is indispensable for effective immune response dynamics, and its dysregulation is critically implicated in the pathogenesis of numerous diseases, such as various cancers and autoimmune disorders [1–5]. During early development, macrophages differentiate into two distinct subtypes based on their origin: tissue-resident macrophages and blood monocyte-derived macrophages (MDMs) [6, 7]. Notably, the precise functional distinctions between tissue-resident macrophages and MDMs remain largely elusive, as both subtypes frequently exhibit overlapping contributions to tissue homeostasis restoration following injury or disease [8–11]. Furthermore, the inherent plasticity of macrophages enables these cells to respond distinctly to a multitude of diverse signals. Macrophage activity is profoundly influenced by external cues, such as cytokines, microbes, Pathogen Associated Molecular Patterns (PAMPs), Damage Associated Molecular Patterns (DAMPs), nucleotide derivatives, antibody-Fc receptor stimulation, glucocorticoids, and stimuli associated with infection and phagocytosis. Given their versatile biology, macrophages represent a highly attractive therapeutic target in drug discovery efforts for both autoimmunity and (immuno-)oncology [12–16].

Human induced Pluripotent Stem Cells (hiPSCs) represent a powerful tool to model development and disease. HiPSCs can differentiate to virtually any cell type in the body and offer key advantages over other model systems which have historically been employed in drug discovery. Specifically, hiPSC-derived cells exhibit enhanced translational relevance compared to traditional immortalised cell lines and, unlike primary cells which often present challenges in large-scale acquisition, hiPSCs provide a scalable *in vitro* system, suitable for high-throughput screening applications. Furthermore, hiPSCs derived from a single individual can be directed towards multiple cellular lineages, facilitating the generation of autologous cell models that more accurately recapitulate complex disease mechanisms. The utility of hiPSCs extends to their application into more sophisticated systems, wherein disease-relevant mutations can be precisely introduced to construct highly translationally relevant cellular models [17–19]. Combined with advancements in gene editing technologies, such as the CRISPR/Cas9 system, hiPSC-derived cell models represent a valuable resource critical for advancing the frontiers of target discovery and drug development.

In this manuscript, we introduce a robust and extensively characterised platform for the large-scale production of hiPSC-derived macrophages, validated using multi-omics and functional assays. We further describe the development and implementation of distinct CRISPR methodologies to conduct genetic screens for drug target identification and validation. Our approach centres on delivering the CRISPR/Cas9 machinery into terminally differentiated hiPSC-derived macrophages, using both arrayed and pooled gene editing formats. We systematically characterise the effects of gene knockouts (KOs) on macrophage biology using a range of phenotypic assays as well as transcriptomics. Our study focuses on two hallmark macrophage stimulations: the combination of LPS and IFN-γ to probe Type 1-skewed phenotypes, and IL-4 to assess Type 2-skewed responses. To the best of our knowledge, this work represents the first parallel application of a high-throughput arrayed CRISPR screening platform and a CRISPR droplet sequencing (CROP-seq) platform to identify critical genetic regulators of hiPSC-derived macrophage polarisation dynamics. These advanced genome editing platforms offer a unique tool for early drug discovery, enabling the identification and validation of novel therapeutic targets by elucidating complex biological mechanisms within macrophages. Through the integration of complementary readouts, we have generated comprehensive data packages for targets discovered in these genetic perturbation screens.

## Results

### Establishment and characterisation of a high-throughput hiPSC-derived macrophage platform

We established a scalable platform for the weekly production of hiPSC-derived myeloid cells, adapted from the protocol by Wilgenburg et al. [20]. Figure 1a outlines the optimised differentiation workflow. Undifferentiated hiPSC cultures at approximately 80% confluency were dissociated into single cells and subsequently incubated in media supplemented with BMP4, VEGF, and SCF within low-attachment 384-well plates to facilitate embryoid body (EB) formation. This step incorporated automated liquid handling to minimise variability and ensure consistent EB size. Four days post seeding, embryoid bodies were transferred to gelatin-coated flasks and cultured in media supplemented with colony-stimulating factor 1 (M-CSF) and IL-3 to promote myeloid lineage specification (week 0). At this stage, the EBs adhered to the gelatinised flasks, a monolayer of cells grew progressively around them and, subsequently, progenitor cells termed “myeloid precursor-like” cells were released into the supernatant (Figure 1a).

**Figure 1.**
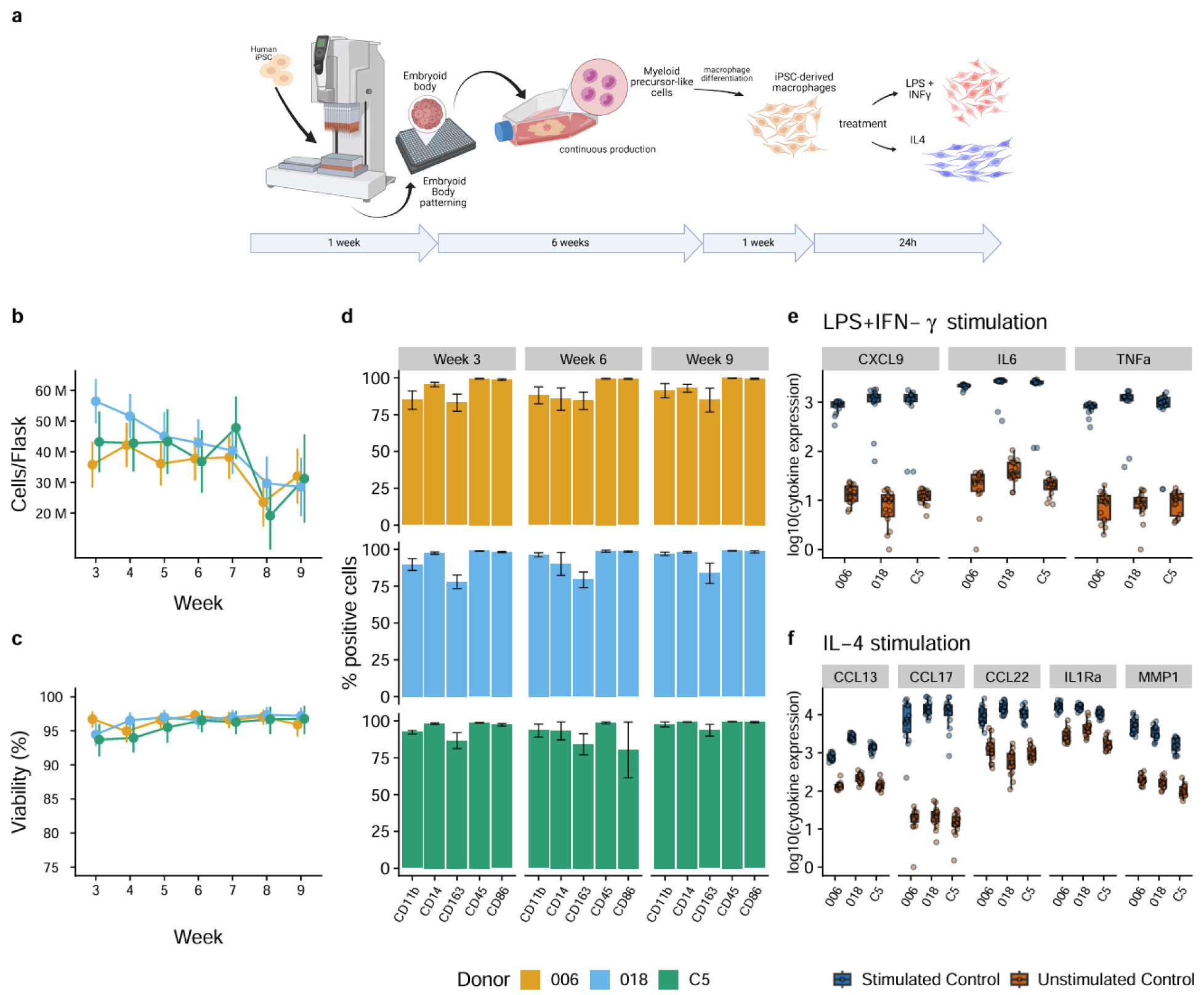
Establishment and characterization of a platform to produce hiPSC-derived macrophages at scale. a) Differentiation diagram showing the timeline from hiPSCs to terminally differentiated, polarised macrophages. Created in BioRender. Carbone, B. (2026) https://BioRender.com/v9e3qdl. b-c) Myeloid precursor-like cell production from week 3 until week 9 was quantified in 3 hiPSC donors. Points show model-estimated marginal means for each donor × week; error bars indicate 95% confidence intervals. d) HiPSC-derived macrophages differentiated from myeloid precursor-like cells harvested on weeks 3, 6, and 9 were analysed for macrophage marker expression by flow cytometry. Expression is quantified as % of positive cells for each marker protein. Bars show mean % positive cells across replicates; error bars represent ± standard error of the mean (SEM). e) Cytokine secretion analysis in cell culture supernatants harvested from day 6 macrophages that had received a 24h stimulation with LPS (100 ng/ml) and IFN-γ (20 ng/ml), or no stimulation as a negative control. f) Cytokine secretion analysis in cell culture supernatants harvested from day 6 macrophages that had received a 24h stimulation with IL-4 (20 ng/ml), or no stimulation as a negative control.

From weeks 3 to 9 during each differentiation experiment, free-floating myeloid precursor-like cells were routinely harvested from the culture media. Weekly production yields of myeloid precursors reached up to 80 million cells per flask, with a gradual reduction to 20 million cells in later weeks, while maintaining consistent viability above 90% (Figures 1b and c). Mature, adherent macrophages were obtained within one week of further differentiation of these myeloid precursor-like cells by plating them on uncoated tissue-culture vessels in medium containing fetal bovine serum and M-CSF (Figure 1d). We assessed the expression of canonical macrophage markers (CD11b, CD14, CD163, CD45, and CD86) in macrophages differentiated from myeloid precursors harvested at weeks 3, 6, and 9. Consistently high expression (>75%) of all markers was observed across multiple hiPSC donors at the protein level (Figure 1d), indicating that the potential of these precursors to generate fully matured macrophages was independent of the culture’s age.

The terminally differentiated hiPSC-derived macrophages demonstrated robust responsiveness to classical stimulations. Treatment with LPS+IFN-γ promoted a Type 1 inflammatory state, while IL-4 treatment promoted a Type 2 inflammatory state. These stimulations represent two defined activation states, while *in vivo* macrophage phenotypes exist along a continuum shaped by complex environmental cues. Macrophage polarisation following LPS+IFN-γ stimulation was evidenced by an increase in the secretion of Type 1 phenotype-associated cytokines, including CXCL9, IL-6, and TNF-α. Conversely, IL-4 treatment induced the upregulation of Type 2 phenotype-associated cytokines, such as CCL13, CCL17, CCL22, IL1Rα, and MMP1 (Figure 1e).

### Multi-omics-based characterisation of hiPSC-derived macrophages

We investigated the cellular and molecular identity of hiPSC-derived macrophages and assessed their similarity to primary monocyte-derived macrophages, henceforth referred to as primary macrophages, using a multi-omics approach. Specifically, hiPSC-derived macrophages and primary macrophages from healthy donors were cultured under identical conditions, stimulated with LPS+IFN-γ or IL-4 or maintained unstimulated, and subsequently analysed using ATAC-seq, RNA-seq, and proteomics (Figure 2).

**Figure 2.**
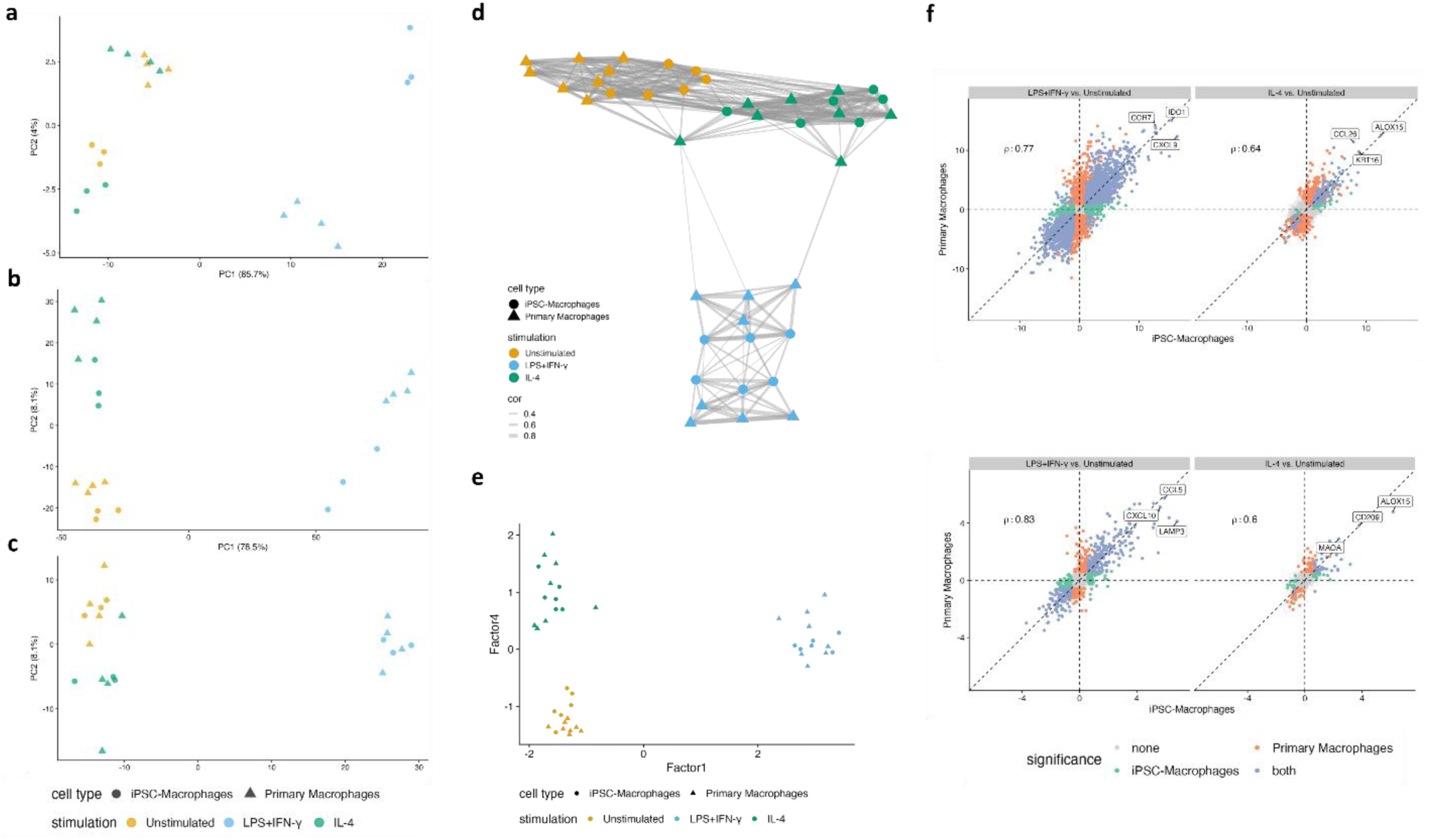
Multi-omics characterization of hiPSC-derived macrophages. a, b, c) PCA plots of (from top to bottom) ATAC-seq, RNA-seq and proteomics analysis of 3 hiPSC-derived and 4 primary macrophage donors either left unstimulated or stimulated with IFN-γ (20 ng/ml) and LPS (100 ng/ml) or with IL-4 (20 ng/ml). d, e) MOFA conducted on hiPSC-derived and primary macrophage omics data in three different polarisation conditions: unstimulated, LPS+IFN-γ, or IL-4. The plot at the top shows separation of the cell types and stimulation states taking into consideration all 10 latent factors analysed. The plot at the bottom was generated using latent factors 1 and 4, which were shown to be critical for the separation of the macrophage stimulation profiles. f, g) Spearman correlation analysis of transcriptomics (top) and proteomics (bottom) data-based differential expression analysis results comparing unstimulated versus LPS+IFN-γ-stimulated or IL-4-treated hiPSC-derived and primary macrophages.

We initially performed principal component analysis (PCA) to explore global omics similarities and stimulation-induced changes across the different cell types and states (Figures 2a, b, and c). We observed that LPS+IFN-γ stimulation constituted the primary driver of PC1, leading to robust changes in cell polarisation, evident at the transcriptomic, proteomic, and epigenetic levels. The effect of IL-4 stimulation appeared less prominent. RNA-seq and proteomics analyses revealed more subtle differences between unstimulated and IL-4-treated cells, with the epigenetic profiles of these two groups clustering closely. To enhance data resolution and investigate more discrete changes, we applied Multi-omics Factor Analysis (MOFA) [21]. Ten latent factors across the three omics datasets were analysed to define the drivers of each stimulation profile across cell types and stimulation conditions. MOFA confirmed the trends observed with individual omics approaches, demonstrating a clear separation of LPS+IFN-γ-stimulated cells from both unstimulated and IL-4-stimulated macrophages, while IL-4-stimulated cells clustered closely with unstimulated samples (Figure 2d). We identified latent factors 1 and 4 as the primary factors driving sample segregation based on stimulation status (Figure 2e). The main features captured by Factor 1 included pathways associated with IFN-γ, IFN-α, and TNF-α signalling, which was expected given the stimulations employed. Latent factor 4, however, revealed a more distinct separation between IL-4-stimulated and unstimulated cells compared to the individual omics analyses. Latent factor 4 involved IL-4, IFN-γ, and IFN-α signalling, as well as cell cycle pathways and E2F transcription factors.

Utilizing the PCA described above to compare hiPSC-derived macrophages against their primary counterparts, and based on the interspersed clustering of the two cell types under both unstimulated and stimulated conditions, it became evident that hiPSC-derived and primary macrophages shared common proteomic profiles (Figure 2c). This finding was largely mirrored by transcriptomics analysis (Figure 2b), with the notable exception of the LPS+IFN-γ condition where primary and hiPSC-derived macrophages exhibited more distinct clustering along the PC2 axis. Interestingly, chromatin accessibility analysis revealed a notable separation of the two cell types across all three conditions (Figure 2a). This observation could be attributed to the distinct ontogeny of hiPSC-derived versus primary macrophages, as described in multiple studies [22].

We also conducted a comprehensive correlation analysis on the results of stimulated-versus-unstimulated cell differential expression analyses, performed on our transcriptomics and proteomics datasets, to compare the overall directionality in gene/protein expression change upon stimulation in hiPSC-derived versus in primary macrophages (Figures 2f and g, Supplementary Figure 1). Notably, high correlation coefficients were observed for both datasets following LPS+IFN-γ exposure, demonstrating a robust and highly similar Type 1 inflammatory response in the two macrophage types. Furthermore, we observed strong agreement in differentially expressed marker profiles in the two cell types, with established pro-inflammatory factors such as CXCL9, IDO1, and CCL5 being among the most significantly upregulated in both hiPSC-derived and primary macrophages. Conversely, IL-4 stimulation yielded lower correlation scores and a reduced number of significantly altered genes/proteins in both macrophage types. Despite this, the top upregulated factors, including ALOX15, CD209, MAOA, and CCL26, all represented established markers of IL-4-mediated responses [23–26], while we show KRT16 to be a novel marker for the IL-4 stimulation condition in hiPSC-derived macrophages.

In summary, our multi-omics approach allowed for extensive characterisation of our hiPSC-derived macrophage model at steady state and upon response to distinct stimulations, regarding features spanning from chromatin remodelling to protein expression. Collectively, our findings demonstrate that hiPSC-derived macrophages exhibit a high degree of similarity to primary macrophages, thereby establishing the suitability of this model for studying human macrophage physiology *in vitro*.

### Development of an RNP nucleofection-based genome editing pipeline in terminally differentiated hiPSC-derived macrophages

CRISPR/Cas9-based functional genomics screens utilising stem cell-derived cellular models are commonly performed at the pluripotent hiPSC or precursor-like cell stage [27–29]. However, this approach presents several limitations, including limited throughput for arrayed screens due to the rapid proliferation of hiPSCs, which prevents their long-term maintenance in 384-well plates, and the potential for gene KOs to impair hiPSC differentiation capacity towards specific lineages. To address these limitations, we developed two distinct workflows for genome editing in terminally differentiated hiPSC-derived macrophages, enabling both arrayed and pooled functional genomics screens.

An arrayed CRISPR/Cas9 ribonucleoprotein (RNP) nucleofection-based genome editing approach was initially optimised for hiPSC-derived macrophages, utilising a pool of three synthetic guide RNAs per gene target. Nucleofection conditions were tested at several time points along the myeloid precursor-like cell-to-macrophage differentiation timeline, and day 5 post-initiation of differentiation was identified as the optimal time point for nucleofection regarding RNP complex delivery efficiency and cell viability. Subsequently, the genome editing pipeline was optimised to determine the most appropriate timing to apply stimulation conditions post-gene knockout. Ensuring the degradation of endogenous proteins prior to assessment of gene knockout effects was critical as part of this optimisation effort, to avoid false results due to residual protein amounts within the cells when evaluating gene knockout phenotypes. To this end, we assessed global proteome turnover in hiPSC-derived macrophages using SILAC (Stable Isotope Labeling by Amino Acids in Cell Culture)-based quantitative proteomics. Six days post-nucleofection, less than 30% residual protein levels were detected, for approximately 80% of the proteome (Figure 3b). Based on these findings, day 11 post-macrophage differentiation initiation was selected as the optimal stimulation timepoint for this protocol.

**Figure 3.**
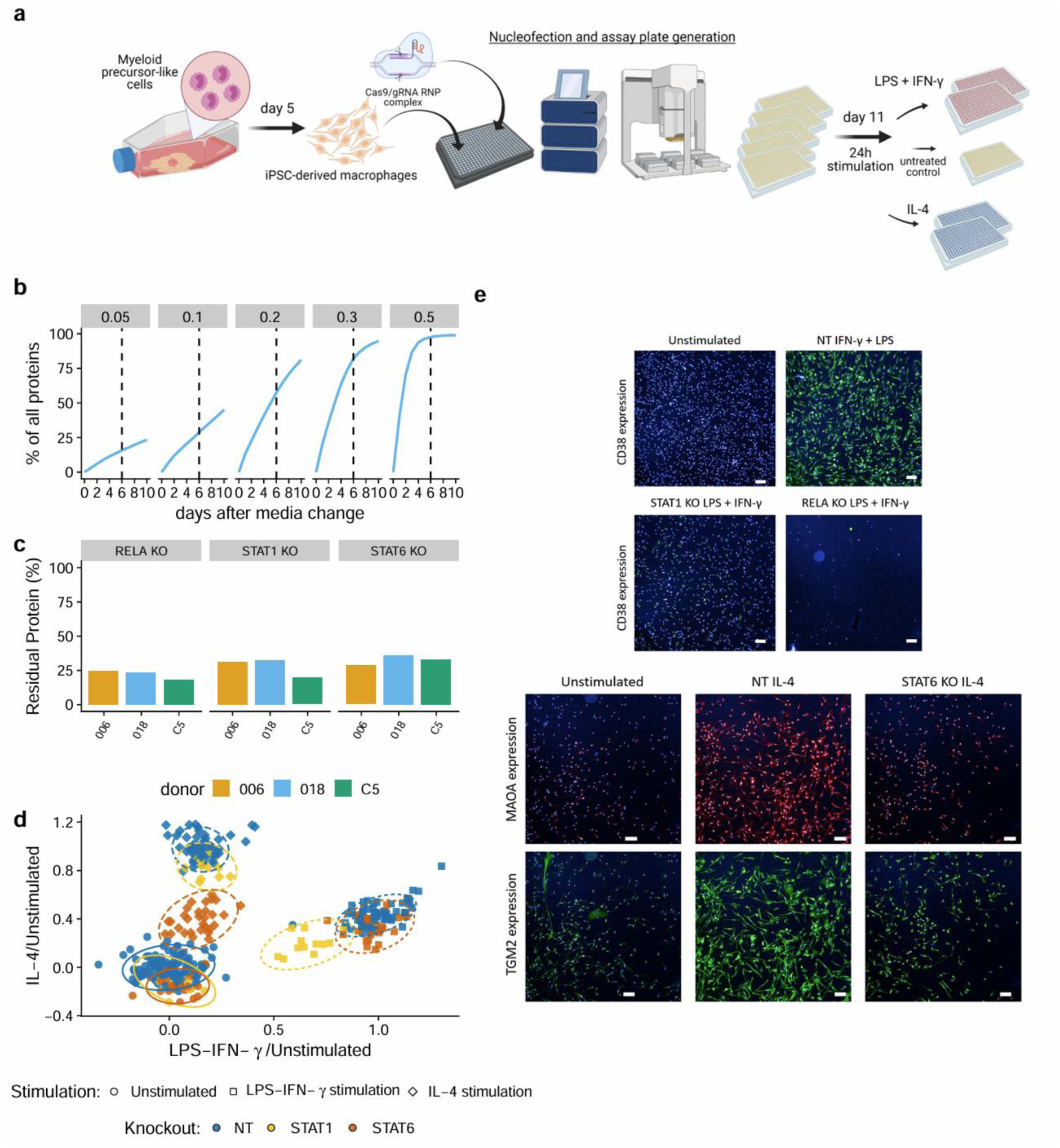
RNP nucleofection-based genome editing pipeline for arrayed KO screening studies in hiPSC-derived macrophages. a) Genome editing is performed on day 5 of the macrophage differentiation protocol. Macrophages are then cultured for an additional 6 days, before being stimulated using either LPS+IFN-γ or IL-4 on day 11, for 24h. Created in BioRender. Carbone, B. (2026) https://BioRender.com/v9e3qdl. b) Dynamic SILAC-based quantitative proteomics study of global proteome turnover in hiPSC-derived macrophages. The facets of the chart represent categories of old protein % present within the cell population from day 0 (day 5 of macrophage differentiation) until day 10 (day 15 of macrophage differentiation). Residual protein percentages are categorized from 0.05% (leftmost facet) to 50% (rightmost facet). c) Genome editing was performed in 3 hiPSC donors to generate KO macrophages for *RELA, STAT1* or *STAT6*. Residual protein levels were measured 6 days post-RNP nucleofection to assess genome editing efficiency. d) EP Analysis of proteomics samples of *STAT1* KO, *STAT6* KO or NT-edited cells, either stimulated with LPS+IFN-γ or IL-4 or left unstimulated. e) Immunofluorescence imaging of genome-edited cells upon stimulation. CD38, MAOA and TGM2 expression was assessed in *STAT1* KO, *STAT6* KO, and *RELA* KO macrophages. White scale bar on each image represents 100μm in length.

To validate our approach, we targeted three known genes implicated in macrophage polarisation biology. *STAT1* and *RELA* are components of the IFN-γ and LPS pathways, respectively, while *STAT6* plays a critical role in IL-4-triggered responses [30, 31]. Following genome editing of these loci, proteomics analysis on day 11 revealed that residual protein levels for all three targets were below 40%, confirming functional KOs for these genes (Figure 3c). To confirm the phenotypic effects of these KOs, we identified markers representative of each stimulation state and assessed their modulation following gene perturbation. Specifically, by interrogating the multi-omics dataset presented above, we identified *CD38* as the top upregulated marker following LPS+IFN-γ stimulation and *MAOA* and *TGM2* as the most highly upregulated factors after IL-4 treatment of hiPSC-derived macrophages, and evaluated their expression via immunofluorescence imaging post-gene KO (Figure 3e). CD38 exhibited robust and homogeneous expression in hiPSC-derived macrophages stimulated with LPS+IFN-γ and transfected with a non-targeting (NT) control gRNA. Upon *STAT1* KO, CD38 expression was undetectable, indicating an attenuation of the LPS+IFN-γ response, as expected. Similarly, MAOA and TGM2 displayed high and consistent levels of expression in IL-4-treated cells transfected with a non-targeting (NT) control gRNA, but their expression was absent in the *STAT6* KO condition, consistent with an anticipated failure of STAT6-deficient macrophages to respond to IL-4 treatment. Unexpectedly, *RELA* KO macrophages exhibited a lethal phenotype upon stimulation with LPS+IFN-γ, which was likely attributable to TNF-α-induced stress [32]. *RELA* was incorporated into our screening workflow as a technical control to assess genome editing and immunostimulation efficiencies prior to downstream analysis.

To obtain a more comprehensive understanding of macrophage pathway responses following gene KO, proteomics analysis was performed on *STAT1* and *STAT6* KO cells following stimulation with LPS+IFN-γ or IL-4, respectively (Figure 3d). The *STAT1* KO LPS+IFN-γ-treated samples distinctly clustered away from the stimulated cell population containing NT gRNAs, shifting towards the unstimulated sample cluster in our plots. Similarly, the *STAT6* KO samples exhibited a comparable effect regarding the corresponding stimulation condition, with the KO cell population shifting away from the IL-4-treated cluster and towards the unstimulated condition cluster. These findings indicated that the selected gene KOs effectively prevented macrophage responses to the corresponding stimulation at the level of the global proteome, rather than merely modulating individual marker expression. These gene KOs were subsequently employed as positive controls for all downstream CRISPR screens. Importantly, these findings were reproduced across multiple hiPSC donors, highlighting the robustness of our workflow.

### An RNP nucleofection-based arrayed CRISPR screen for macrophage polarisation-associated target discovery at scale

We generated two gene libraries as target input for genetic screening using our arrayed genome editing pipeline in hiPSC-derived macrophages stimulated with LPS+IFN-γ or IL-4, drawing from multi-omics-based expression data as well as established signalling pathways relevant to macrophage biology. These libraries contained 258 and 262 gene targets for screening in LPS+IFN-γ-stimulated or IL-4-treated macrophages, respectively, and partially overlapped (Figure 4a). To enable high-throughput arrayed CRISPR screening, we adapted our gene editing protocol to a 384-well format, establishing an automated workflow that generated 12 replicate assay plates from a single nucleofection plate. The macrophages within the replicate assay plates were then stimulated, and parallel downstream assays provided multiparametric readouts for our genetic screens (Figure 4a). This comprehensive screening effort involved 1.2 billion macrophages derived from three different hiPSC donors, and was executed in two distinct arms. For the LPS+IFN-γ arm, screen readouts included assessing CD38 upregulation by immunofluorescence imaging coupled to ELISA-based quantification of Type 1 cytokine/chemokine (CXCL9, IL6, and TNF-α) secretion levels. Similarly, the IL-4 arm involved profiling MAOA and TGM2 upregulation by immunofluorescence microscopy alongside ELISA-based quantification of an expanded panel of Type 2 cytokines (CCL13, CCL22, CCL24, IL1Rα, and MMP1).

**Figure 4.**
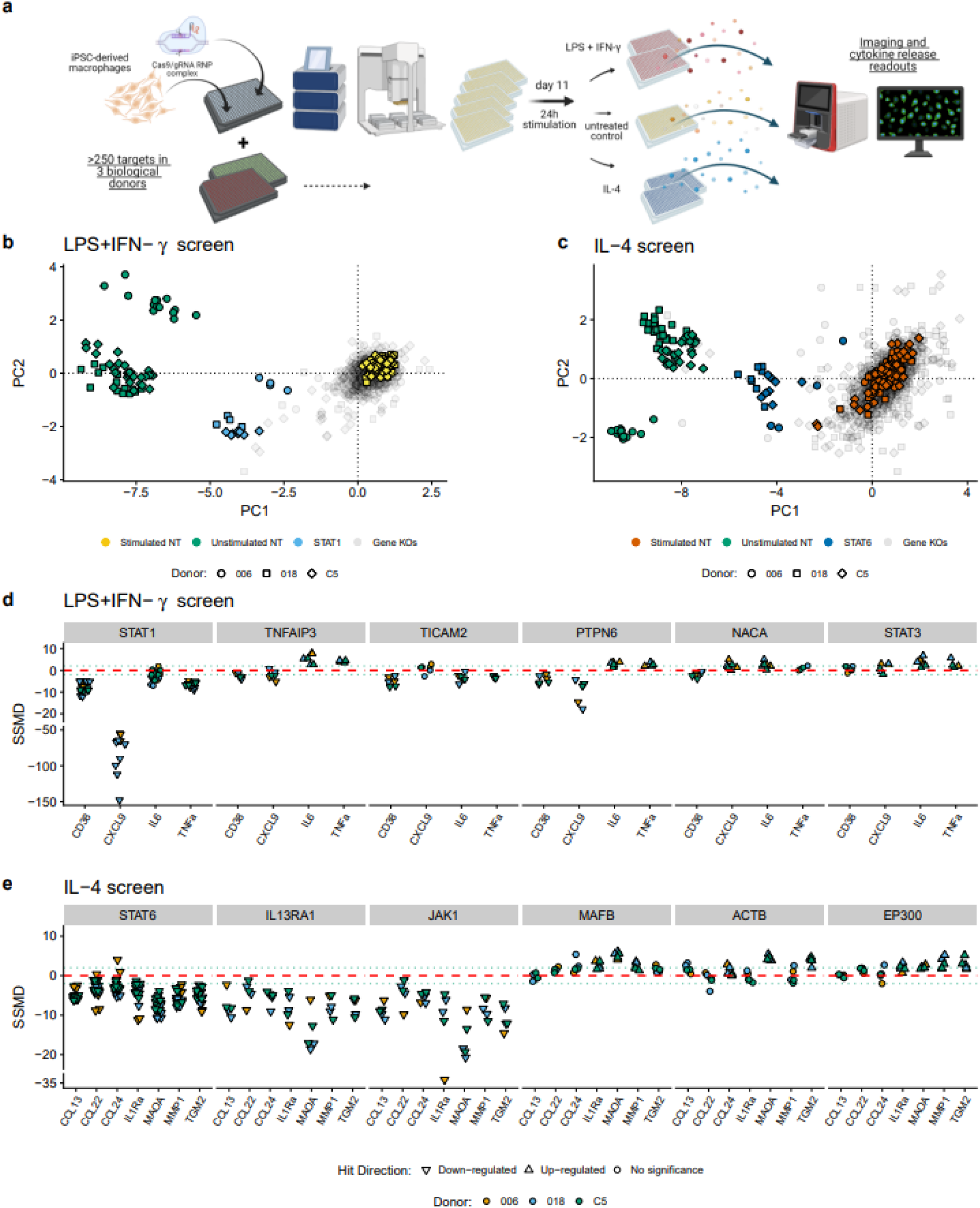
Arrayed CRISPR KO screening for macrophage polarisation by LPS+IFN-γ or IL-4. a) Schematic illustrating the experimental workflow followed during execution of two arrayed CRISPR KO screens in hiPSC-macrophages polarised using two different stimulation conditions: LPS+IFN-γ or IL-4. Both genetic perturbation screens were performed on 3 different hiPSC donors. In this workflow, genome editing was conducted using an automated protocol allowing for the generation of 12 assay plates from a single electroporation plate. Those assay plates were kept in culture until day 11 of this workflow (an additional 6 days post-nucleofection). On day 11 of the workflow, the gene-edited macrophages within the assay plates were stimulated with either LPS+IFN-γ or IL-4, for 24h. The next day, each assay plate was used for cytokine secretion and high content imaging readouts. Schematic generated using BioRender. Carbone, B. (2026) https://BioRender.com/v9e3qdl. b and c) PCA plots depicting the cytokine release and high content imaging assay results acquired for the LPS+IFN-γ and IL-4 KO screens in hiPSC macrophages. Control conditions (macrophages nucleofected with non-targeting gRNAs and subsequently cultured with or without stimulation, as well as the STAT1 or STAT6 KO controls for the LPS+IFN-γ and IL-4 KO screens, respectively) have been labelled in colour, whereas all gene KOs evaluated as test conditions during the two screens are labelled in grey in our PCA plots. d and e) Selected examples of gene KOs included in the LPS+IFN-γ or IL-4 CRISPR screens in hiPSC-macrophages and effects of each gene KO on the expression/secretion levels of the marker panels included in each readout assay employed for each screen. Results reported using SSMD (Strictly Standardized Mean Difference). Green horizontal lines represent the designated SSMD cutoff of ±2.

Principal component analysis (PCA) was performed on the results of the aforementioned assays for each of our two CRISPR screens in hiPSC-derived macrophages. Successful genome editing and macrophage polarization were evidenced by the shift of the *STAT1* and *STAT6* KO clusters towards the unstimulated macrophage clusters in our plots (Figures 4b and c). This analysis revealed a spectrum of gene KO effect strengths on functional macrophage phenotypes, as illustrated by gene KO dispersion across our PCA plots. Gene KO effects were quantified using the Strictly Standardized Mean Difference (SSMD) [33], a statistical measure of effect size and directionality commonly employed in high-throughput screening for hit identification. SSMD was calculated for each endpoint by comparing the signal from the stimulated gene KO sample to the non-targeting (NT) control within the same stimulation condition, thereby quantifying the specific KO effect on the cellular phenotypes under investigation. Hit calling criteria for target gene identification were adjusted based on the number of readouts per screen. For the LPS+IFN-γ screening arm, genes with an SSMD of ±2 in at least two endpoints across a minimum of two hiPSC donors were classified as hits, resulting in 44 genes being identified. For the IL-4 screen, which involved a greater number of readouts, hit calling was based on gene KOs exhibiting upregulation or downregulation in at least three endpoints with an SSMD of ±2 in a minimum of two hiPSC donors, leading to the identification of 19 gene hits.

Gene hits with strong SSMD values, representative of the marker modulation effects exerted by their KO, are presented in Figures 4d and e for the LPS+IFN-γ and IL-4 screens, respectively. While some gene KOs consistently triggered either downregulation or upregulation of all markers measured, suggesting broad suppression or enhancement of macrophage activation pathways, others, such as *TNFAIP3* and *PTPN6* KOs, exhibited a mixed phenotype, characterised by upregulation of IL-6 and TNF-α coupled with downregulation of CXCL9 and CD38 (Figures 4d and e). Interestingly, mixed phenotypes were predominantly observed in the LPS+IFN-γ screening arm, suggesting that macrophage activation in this condition is driven by pathway-specific rewiring rather than strictly binary activation states. In the IL-4 screen, gene KOs such as *IL13RA1* and *JAK1* KOs led to strong marker downregulation, while others including *MAFB*, *ACTB*, and *EP300* KOs resulted in robust marker upregulation in our readouts. These findings showcase the sensitivity of our CRISPR screening workflow in capturing expected as well as novel and strong as well as subtle biological effects associated with macrophage polarisation under two distinct stimulatory conditions.

To our knowledge, this study represents the first multiparametric, high-throughput arrayed CRISPR knockout screen conducted in terminally differentiated hiPSC-derived macrophages. The scale of our genome editing workflow allows for rapid and precise perturbation of hundreds of genes simultaneously, significantly expanding the search space for novel therapeutic targets. Further downstream target prioritisation efforts can integrate pathway relevance, target tractability, and consistency across readouts as decision-making criteria. By providing multi-dimensional, context-specific functional insights, such studies enhance our understanding of fundamental disease mechanisms, aid in de-risking candidates, and ultimately allow for streamlining the drug development pipeline with a high degree of translational relevance.

### Development of a CROP-seq method for pooled CRISPR screening in terminally differentiated hiPSC-derived macrophages

Our arrayed CRISPR screening platform relies on established assays, potentially narrowing its application to disease biology for which development of functional assays is already established or generally feasible and rendering it less applicable to more nascent biological questions where a more agnostic, exploratory approach is needed. Pooled CRISPR screening methods such as CROP-seq (pooled CRISPR screening followed by single-cell RNA sequencing) overcome this limitation by providing an unbiased transcriptomics readout adapted for screening for novel biology at single-cell resolution. Such comprehensive molecular profiling, directly coupled with individual genetic perturbations, is particularly advantageous for exploring novel disease areas or cellular processes for which robust, high-throughput functional assays have not yet been developed.

We optimised a CRISPR droplet sequencing (CROP-seq) method tailored to terminally differentiated hiPSC-derived macrophages. Our strategy builds upon the Single guide RNA (sgRNA) Lentiviral Infection with Cas9 protein Electroporation (SLICE) protocol, incorporating lentiviral infection of cells with a modified CROP-seq gRNA vector, subsequently followed by Cas9 protein electroporation [34]. Our approach is similar to the SLICE protocol, including infection of cells with a lentivirus carrying the modified CROP-seq gRNA vector, followed by Cas9 protein electroporation [34–36]. To ensure that each cellular phenotype observed could confidently be attributed to a singular gene perturbation, we employed a low Multiplicity Of Infection (MOI) of approximately 0.3 in our experiments. The experimental timeline involved lentiviral transduction on day 5 (post-initiation of differentiation of myeloid precursor-like cells to macrophages), Cas9 protein electroporation on day 7, and a subsequent cell sorting step on day 12 to enrich for gRNA-expressing cells. Following recovery until day 14, cells were stimulated with LPS+IFN-γ. Twenty-four hours after stimulation, macrophages were harvested and subjected to single-cell transcriptomics with gRNA capture analysis using the 10x Genomics Chromium Platform (Figure 5a).

**Figure 5.**
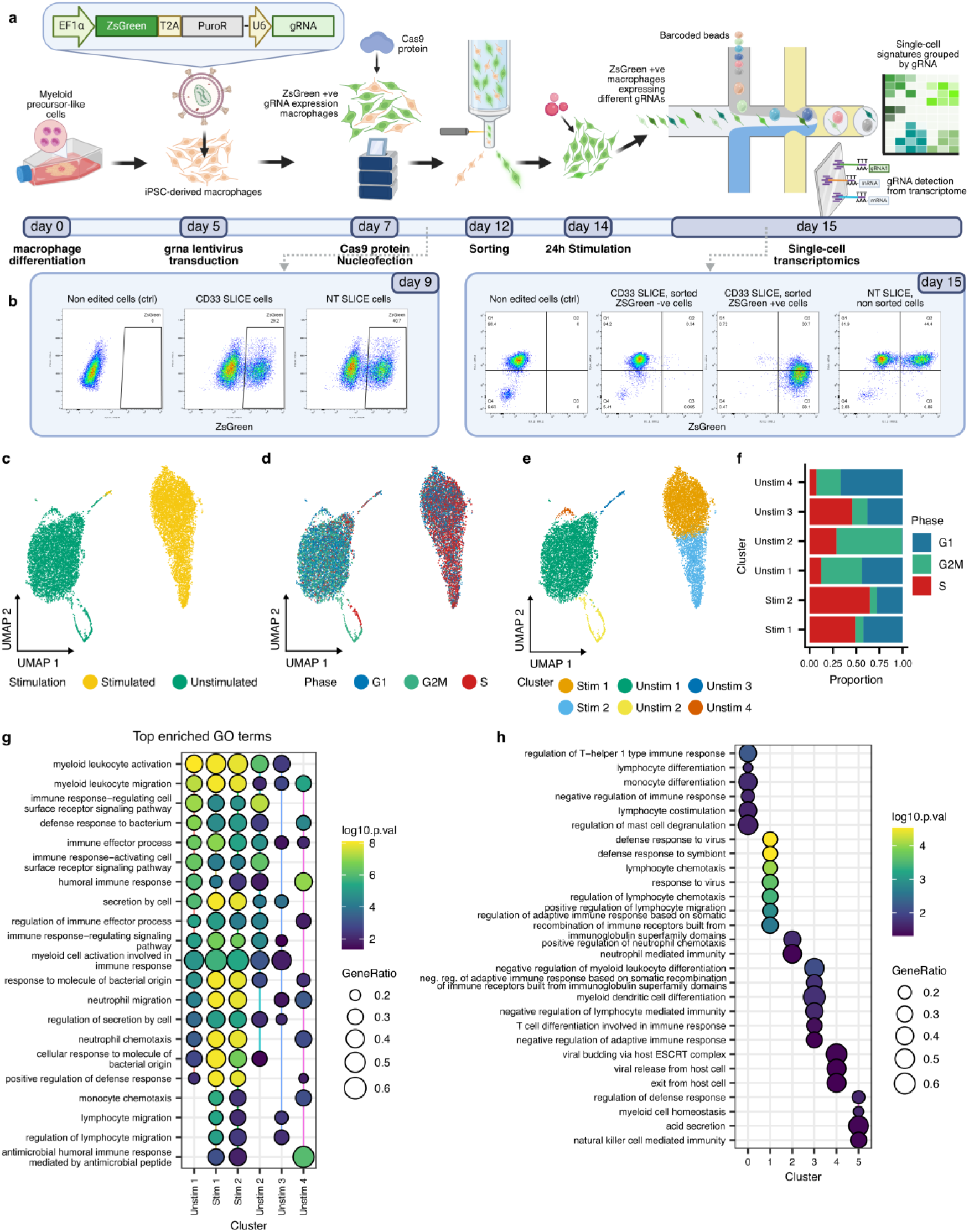
Development of a CROP-seq method tailored to hiPSC-derived macrophages and execution of a pooled screen. a) Schematic outlining our CROP-seq method coupled to SLICE, adapted for terminally differentiated hiPSC-derived macrophages. As part of this workflow, myeloid precursor-like cells were differentiated towards macrophages for 5 days. On day 5, macrophages were transduced with a lentivirus expressing a modified CROP-seq gRNA construct. Transduced macrophages were then electroporated with Cas9 protein on day 7, and sorted based on ZsGreen expression on day 12. Cells were subjected to LPS+IFN-γ stimulation on day 14. After 24h, cells were processed for single-cell transcriptomics with gRNA capture analysis. Created in BioRender. Carbone, B. (2026) https://BioRender.com/v9e3qdl. b) Flow cytometry-based analysis of macrophages processed using the SLICE protocol, showing a comparison between a CD33 KO and non-targeting gRNA control condition at the protein level. c and d) UMAP analysis of single-cell RNA-seq data acquired for WT macrophages, depicting clustering patterns occupied by stimulated versus unstimulated cells as well as distinct cell cycle phases. e) UMAP plot depicting subclusters identified for the stimulated and unstimulated WT macrophage populations, identified through unsupervised clustering analysis of the WT macrophage single-cell transcriptomics dataset. f) Enrichment of the newly defined stimulated and unstimulated WT macrophage subclusters for the different phases of the cell cycle. g and h) Top enriched GO terms for each of the newly identified stimulated and unstimulated WT macrophage clusters.

Lentiviral transduction in macrophages is notoriously challenging, often requiring the use of specialised tools such as Vpx constructs or SAMHD1 depletion to overcome antiviral mechanisms endogenous to this immune cell type [29, 37, 38]. Our CRISPR screening strategy achieved high genome editing efficiencies without the need to incorporate such tools into the workflow. For example, analysis by flow cytometry revealed that our method achieved successful protein-level KO in ∼70% of the transduced macrophage population for CD33, which we employed as a technical control (Supplementary Figure 2c). Our workflow also demonstrated no detrimental impact on the ability of the cells to respond to immunostimulation, as evidenced by cytokine release analysis of gene-edited macrophages after treatment with LPS+IFN-γ or IL-4 (Supplementary Figures 2a and b).

We executed a pooled CROP-seq screen in macrophages derived from three hiPSC donors, targeting 16 genes that demonstrated robust KO phenotypes in our arrayed LPS+IFN-γ screen described above. This list included candidates that elicited either an enhancement or a suppression of the cellular response to LPS+IFN-γ stimulation as previously assessed by our arrayed screening readouts. The pooled library contained 4 gRNAs per target gene, designed using the CRISPick software [39], as well as 8 gRNAs targeting the AAVS1 safe harbour locus and 8 non-targeting gRNAs, resulting in a total panel of 80 gRNAs [40]. Genome editing efficiency was validated at the protein level via a parallel flow cytometry assay using CD33 KO as a control (Figure 5b).

To control for potential effects introduced by the sequential manipulation steps inherent to our workflow, we included non-transduced, non-genome-edited macrophage samples, henceforth referred to as wild type (WT) cells, into our screen. We initially focused the analysis of our CROP-seq screening data on this WT macrophage population, to establish a benchmark for non-manipulated cell single-cell transcriptome profiles and enable accurate downstream comparisons to our genome-edited samples. Consistent with our prior bulk transcriptomics dataset on hiPSC-derived macrophages (Figure 2), unstimulated and LPS+IFN-γ-stimulated cells formed distinct clusters in our UMAP plots (Figure 5c). Further analysis revealed an intriguing clustering of cell populations based on their cell cycle stage (Figure 5d), with stimulated cells predominantly occupying G1 or S cell cycle phases and unstimulated cells mainly found in G1 or G2/M phases. Unsupervised clustering applied to the transcriptome profiles of the unstimulated cells further delineated four distinct groups, characterised by one predominant transcriptomic signature and three minor surrounding clusters. Conversely, stimulated samples resolved into two discernible subpopulations, one notably enriched for pathways associated with cellular responses to pathogens and lymphocyte chemotaxis, and the other for neutrophil-mediated immunity and neutrophil chemotaxis pathways (Figures 5e-h). Considering the cell cycle heterogeneity we observed within the unstimulated cell population and recent evidence on the impact of the cell cycle on macrophage polarisation dynamics [41], the emergence of these two M1-like subpopulations in our data might reflect differential responses to IFN-γ and LPS signalling influenced by intrinsic cellular states such as distinct phases of the cell cycle.

Subsequently, gRNA capture was confirmed and analysed for the transduced macrophage population (Figure 6a, Supplementary Figure 3). Within this cell population, 60% to 90% of macrophages exhibited single or multiple gRNA assignments. Aiming to analyse individual gene knockout effects, we exclusively considered cells with single gRNA assignments for our analysis, which accounted for over 40% of all cells across our samples (Figure 6a, Supplementary Figure 3). Importantly, the number of cells identified per gRNA closely correlated with gRNA counts within our lentiviral library, indicating successful maintenance of gRNA representation throughout the SLICE workflow. However, we observed that certain gRNAs were under-represented in our data in terms of cell counts. For instance, unstimulated cells harbouring gRNAs targeting *RELA* were present at only 63% of the anticipated level based on a linear model fitted through the origin (Supplementary Figure 3h). *RELA* gRNA under-representation was exacerbated in stimulated macrophages, where gRNA-carrying cell numbers reached only 34% of the expected value. This observation likely reflects the essential role of *RELA* in macrophage survival, particularly upon stimulation with LPS+IFN-γ, where *RELA* KO would lead to cell death (Figure 3e).

**Figure 6.**
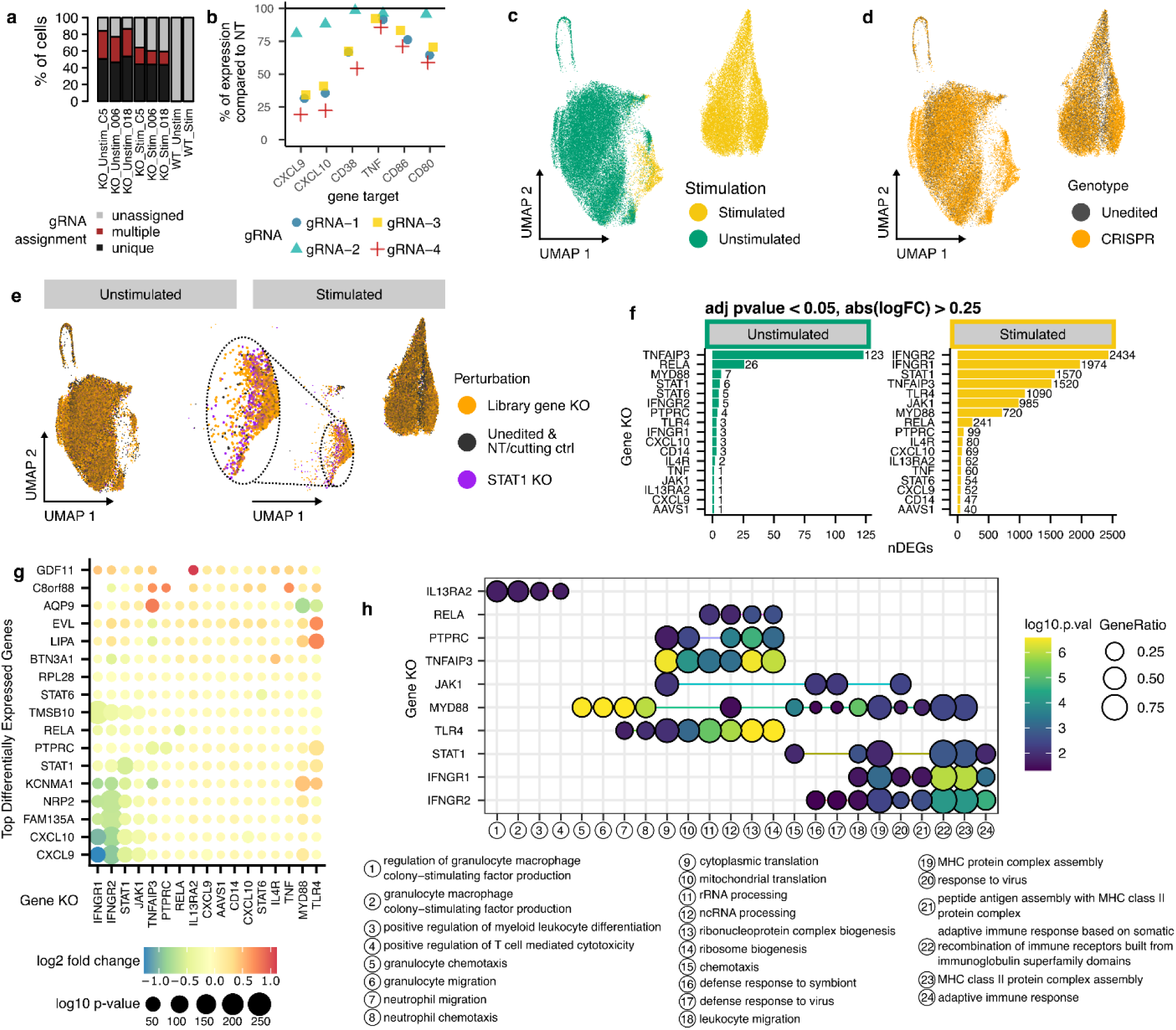
Characterisation of the effects of gene perturbations on the transcriptome of unstimulated and LPS+IFN-γ-treated hiPSC-derived macrophages at single cell resolution. a) GRNA assignment in unstimulated and stimulated macrophages derived from 3 hiPSC donors. The graph shows the percentage of cells with one or more gRNAs or with a lack of assigned gRNAs. b) Effect of *STAT1* gRNAs on STAT1 target gene mRNA expression levels, reporting % of target gene expression in relation to each individual *STAT1* gRNA. c) UMAP analysis plot depicting our CROP-seq data, labelled for the LPS+IFN-γ-stimulated and unstimulated macrophage populations. d) UMAP analysis plot showing our CROP-seq data, labelled for the genome-edited and unedited macrophage populations. e) UMAP analysis plot depicting our CROP-seq data, labelled for cluster enrichment with our library gene-targeting gRNAs, *STAT1* gRNAs, and non-targeting/cutting control gRNAs as well as unedited cells. Stimulated *STAT1* KO macrophages are zoomed in to highlight the unique clustering pattern of this control condition. f) Target gene differential expression assigned to and quantified for each gene KO present in our lentiviral library, relative to the NT control condition. g) Top differentially expressed target genes corresponding to each gene perturbation, in LPS+IFN-γ-stimulated macrophages. h) Gene Ontology term enrichment analysis for the top differentially regulated genes assigned to each gene KO present in our CROP-seq screen, in LPS+IFN-γ-polarised macrophages.

To validate the impact of our gene perturbations on their respective target genes, we explored the effects of *STAT1* KO on downstream STAT1 targets, similar to our arrayed LPS+IFN-γ screen where we employed *STAT1* KO as a known biological control. We quantified the transcript abundance levels of several established STAT1 target genes, including *CXCL9*, *CXCL1*, *CD38*, *TNF*, *CD86*, and *CD80* (Figure 6b). Notably, we observed a substantial downregulation of *CXCL9* and *CXCL10* as a result of *STAT1* KO across all gRNAs, with more modest effects on the remaining targets within the gene panel we examined. These findings confirm the efficacy of our method in detecting transcriptome-level alterations resulting from specific gene perturbations.

UMAP analysis of our CROP-seq data demonstrated clear segregation of the stimulated from the unstimulated macrophage samples, with a distinct, minor stimulated cell cluster observed in close proximity to the primary unstimulated cell population (Figure 6c). Labelling the samples on our UMAP plots according to their corresponding gene perturbation revealed that macrophages receiving non-targeting or AAVS1 gRNAs were uniformly distributed across both the stimulated and unstimulated cell clusters (Figure 6d, Supplementary Figure 3j). This finding affirms the absence of a discernible cellular phenotype attributable to the genome editing protocol itself. Notably, the profound effects of *STAT1* KO on macrophage function appeared as a distinct stimulated cell cluster situated proximally to the unstimulated macrophage population in our UMAP plots. This showed that LPS+IFN-γ-stimulated cells harbouring *STAT1* gRNAs exhibited transcriptomic profiles which closely resembled an unstimulated macrophage state (Figure 6e). This observation is consistent with our arrayed LPS+IFN-γ screen findings and substantiates that, in the absence of STAT1, macrophages fail to establish a functional pro-inflammatory-like phenotype, an effect detectable at the single-cell transcriptome level.

Next, we explored the effects of all 17 gene KOs on differential gene expression in hiPSC-derived macrophages at baseline and upon LPS+IFN-γ stimulation. We observed minimal impact of most gene perturbations on the transcriptome of unstimulated cells, with only *TNFAIP3* and *RELA* KOs modulating the expression of 123 and 26 targets, respectively (Figure 6f). In stark contrast, stimulated macrophages exhibited a significant number of Differentially Expressed Genes (DEGs) as a result of specific gene KOs. In more detail, *IFNGR2*, *IFNGR1*, *STAT1*, and *TNFAIP3* KOs each led to differential expression of over 1500 genes, while *TLR4* and *JAK1* KOs caused differential regulation of more than 900 targets (Figure 6f). Examination of individual target transcripts (Figure 6g) further elucidated these genetic perturbation effects: *IFNGR2*, *IFNGR1*, *STAT1*, and *JAK1* KOs consistently downregulated pro-inflammatory markers crucial for viral infection, such as *CXCL9*, *CXCL10*, *FAN135A*, and *NRP2*. Conversely, *TLR4* KO led to the upregulation of *EVL* and *LIPA*, which are implicated in suppressing cell migration and regulating the lipid-associated pro-inflammatory macrophage state [42].

Gene Ontology (GO) term enrichment analysis corroborated the findings described above, demonstrating that *IFNGR2*, *IFNGR1*, *STAT1*, and *JAK1* KOs significantly impacted chemotaxis, viral immune response, and adaptive immunity associated pathways. In contrast, the effects of *TLR4* KO were more centred on intracellular functions, including ribosomal and mitochondrial processes, as well as neutrophil chemotaxis pathways (Figure 6h).

Collectively, these results confirm the robustness of our CROP-seq workflow in generating precise gene perturbations and analysing their phenotypic effects at the transcriptome level in hiPSC-derived macrophages. Our method effectively captures the effects of gene KOs on disease-relevant pathways and processes at single-cell resolution without compromising cellular physiology, enabling the identification of canonical as well as non-canonical gene regulators of macrophage biology. Importantly, our workflow represents a highly scalable platform for pooled CRISPR screening, providing a powerful tool for drug target discovery efforts. To our knowledge, our study represents the first instance of a pooled CROP-seq screening method towards identification of genetic regulators of macrophage polarisation in terminally differentiated hiPSC-derived macrophages.

## Discussion

Here, we showcase the value and potential of hiPSC-derived models for use in drug discovery. First, we describe an optimised, scalable protocol for hiPSC-to-macrophage differentiation, and present a comprehensive multi-omics characterisation of this hiPSC-derived macrophage model under both pro-inflammatory (LPS+IFN-γ) and anti-inflammatory (IL-4) stimulation conditions. Second, we introduce two distinct, complementary high-throughput CRISPR screening platforms — arrayed and pooled — optimised specifically for hiPSC-derived macrophages. We employ these perturbation screening platforms to uncover key genetic regulators of macrophage polarisation after exposure to pro-inflammatory (LPS+IFN-γ) or anti-inflammatory (IL-4) stimuli. To our knowledge, our study represents the first application of both arrayed and pooled CRISPR screening workflows leveraging diverse readouts, including functional assays and single-cell transcriptomics, to terminally differentiated hiPSC-derived macrophages. Combined with the inherent scalability of hiPSC-derived models, these advanced genetic screening approaches constitute powerful tools for high-throughput functional genomics and drug target identification efforts in human macrophages.

Our scalable hiPSC-to-macrophage differentiation platform is an evolution of existing methodologies for large-scale macrophage production from hiPSCs [43–45]. Moreover, our integrated multi-omics data on the hiPSC-macrophage model, encompassing chromatin accessibility, gene/protein expression, and functional phenotypes such as cytokine secretion, revealed a high degree of similarity between hiPSC-derived macrophages and primary macrophages, across all conditions tested. While hiPSC-derived macrophages are ontogenetically distinct from blood monocyte-derived macrophages, exhibiting features of a tissue-resident precursor-like identity and some functional divergence [46, 47], the marked similarity between hiPSC-derived and primary cells evident in our multi-omics analysis validates the utility of hiPSC-derived macrophages as a physiologically relevant model. Our comprehensive multi-omics data crucially complement prior model characterisation efforts and offer a valuable resource across disease areas with known macrophage involvement, such as in various cancers where tumor-associated macrophages play key roles in immune suppression [46, 48, 49].

Previously published high-throughput CRISPR screens in human macrophages have predominantly utilised macrophages derived from the THP-1 monocytic cell line as the model of choice [50, 51]. These studies employed diverse readouts to uncover genes regulating various immune mechanisms and pathways, including macrophage polarisation in response to LPS, phagocytosis, and pathogen interactions [51, 52]. While these THP-1 screens have significantly advanced CRISPR/Cas9 technology development in human macrophages and provided valuable insights into macrophage biology, one of their key limitations is the reduced translational relevance of the THP-1 cancer cell line compared to hiPSC-derived macrophages as a human macrophage model [51].

Our arrayed CRISPR screening platform for hiPSC-derived macrophages, based on RNP nucleofection, integrates diverse readouts to yield multiparametric data from single gene-edited cell populations, enabling simultaneous interrogation of multiple biological pathways and phenotypic responses. Such readouts include functional assays such as immunofluorescence imaging and cytokine secretion analysis, as presented here, but can also expand to miniaturised omics assays to profile thousands of genes/proteins per gene perturbation.

Our pooled CROP-seq platform employs a variation of the SLICE protocol [36], leveraging single-cell transcriptomics to explore the effects of individual gene KOs in hiPSC-macrophages. Notably, this CROP-seq approach enables dissection of the phenotypes of heterogeneous cell populations, such as different macrophage polarisation states, yielding comprehensive and unbiased datasets particularly valuable for disease biology areas lacking foundational data and functional assays. While we have presented a small-scale CROP-seq study on immunostimulated versus unstimulated hiPSC-derived macrophages, this approach is highly scalable, even towards genome-wide CRISPR screening.

Interestingly, while Vpx constructs are commonly employed to facilitate lentiviral transduction and stable transgene expression in macrophages [29, 37, 38], we found usage of such tools to not be necessary in our study. We hypothesise this is attributable to a combination of our vector’s small size as well as high viral concentration and purity. Additionally, the low proliferation rate of hiPSC-derived macrophages during transduction likely facilitates lentiviral integration. This is supported by evidence that SAMHD1 undergoes phosphorylation by cell cycle-dependent kinases during the S-phase. This phosphorylation event is known to modulate SAMHD1’s dNTPase activity, thereby ensuring sufficient dNTP levels for DNA replication and concurrently reducing SAMHD1-mediated restriction of lentiviral reverse transcription and integration, thus creating a more permissive environment for gRNA delivery in our model system [54].

While a previously published study reported a pooled screening protocol entailing delivery of a genome-wide CRISPR/Cas9 lentiviral library into hiPSC-derived macrophage precursors [29], our distinct method achieves pooled genome editing in terminally differentiated hiPSC-derived macrophages. Importantly, our protocol incorporates a modified CROP-seq vector, enabling a single-cell RNA-sequencing readout while retaining compatibility with other assays such as flow cytometry. A similar pooled CRISPR screening approach employing a modified CROP-seq vector was recently applied to human primary macrophages by Fandrey *et al.*, but with an optical readout [38]. Moreover, while this manuscript was in preparation, Traxler and colleagues independently presented a CROP-seq screen in a murine macrophage cell line [53]. Compared to the Traxler *et al.* study, our workflow avoids reliance on constitutive Cas9 expression and therefore bypasses the time and effort it takes to generate hiPSC lines engineered to express Cas9, instead employing electroporation of Cas9 protein into the cells.

As mentioned above, a key advantage of our two CRISPR screening workflows lies in performing genome editing on terminally differentiated macrophages, rather than at the pluripotent hiPSC or multipotent myeloid progenitor stage. Crucially, genetic modification at earlier developmental stages risks interfering with critical stem/progenitor cell differentiation pathways, potentially yielding aberrant functional responses and false results in such CRISPR screening approaches. However, leveraging earlier stages within our hiPSC-derived myeloid cell differentiation workflow could also provide unique advantages for addressing specific biological questions. For example, beyond the terminally differentiated macrophages central to this study, the myeloid progenitor intermediates produced as part of our adapted differentiation protocol can also be directed towards microglia, the brain’s resident macrophages. Human iPSC-derived microglia constitute a particularly attractive *in vitro* model for basic research and drug discovery, given the extreme difficulty and scarcity in obtaining primary, brain-resident microglia [55]. Consequently, scalable hiPSC differentiation to microglia and high-throughput genetic perturbation screening in terminally differentiated hiPSC-derived microglia represent crucial tools for studying and discovering novel therapeutics for neurodegenerative diseases [56]. Mirroring our dual-platform strategy, both RNP-based arrayed KO and lentivirus-based pooled CRISPRi/a screening methods have been applied to hiPSC-derived microglia to identify modifiers of lipid droplet formation and genes governing microglia survival and function (specifically activation and phagocytosis), showcasing the power in coupling physiologically relevant hiPSC-derived myeloid cell models with CRISPR screening for target discovery [57–59]. Lastly, gene perturbation screening in myeloid progenitors (i.e., monocyte-like cells) can be employed to investigate homeostasis- and disease-relevant processes such as monocyte differentiation to macrophages or tissue infiltration. Collectively, the versatility of our modular hiPSC-to-myeloid-cell differentiation and CRISPR screening platforms significantly expands the available toolkit for dissecting the multifaceted roles of myeloid cells in human disease biology.

Here, we employed our two CRISPR screening methods to genetically dissect the response of hiPSC-derived macrophages to stimulation with LPS+IFN-γ or IL-4, established paradigms for Type 1 and Type 2 macrophage inflammatory states, respectively [60]. However, to comprehensively understand the complex roles myeloid cells play in human disease mechanisms, utilisation of more disease-specific stimuli is essential. Recent literature has highlighted novel stimulation cocktails, such as TNF, Prostaglandin E2, and the TLR1/2 agonist Pam3CSK4, which were shown to induce a chronic inflammation phenotype in macrophages [61, 62]. Furthermore, to holistically address the need for novel, disease-specific macrophage stimulations, we and others have implemented AI/ML-based approaches that integrate multi-omics patient datasets to predict factors inducing bespoke disease signatures *in vitro* [63, 64]. Acknowledging the high attrition rate in drug development programs [65], often stemming from initial target identification in non-physiologically relevant models that poorly recapitulate disease physiology [66], we believe that our genome editing pipelines, integrated with disease-specific macrophage stimulations, will enhance the quality of drug targets identified via functional genomics screening in myeloid cell types. The comprehensive macrophage multi-omics datasets and CRISPR screening platforms generated herein can further support the streamlining and acceleration of drug discovery strategies by enabling omics signatures and functional assays as critical decision points for target progression in therapeutic areas with unmet clinical need.

## Methods

### HiPSC culture

Human induced Pluripotent Stem Cell lines 66540088 (WTSli018A), 66540004 (UKBi006A), 66540188 (WTSli075A) were obtained from the European Bank for Induced Pluripotent Stem Cells (EBiSC) and the samples received Ethics Committee approval (CEI-117-2357) from the originating institutions. The EBiSC bank acknowledges Wellcome Sanger Institute being the source of the lines 66540088 and 66540188, and Universitätsklinikum Bonn being the source of the line 66540004. These lines were generated with support from the Innovative Medicine Initiative (IMI) Joint Undertaking (JU) captured under grant agreement n°115582 and from the IMI-2 JU under grant agreement n°821362. These resources include financial contribution from the European Union’s Seventh Framework Programme (FP7/2007-2013), European Union’s Horizon 2020 research and innovation programme and EFPIA. Human induced Pluripotent Stem cell line iPSCORE_1_13 (S02315_C5) was obtained from the Frazer lab at the University of California San Diego (USCD). All donors provided written informed consent for use of these lines. The lines were sourced ethically, and their research use was in accordance with the terms of the informed consents under an IRB/REC approved protocol.

Cells were cultured on vitronectin (STEMCELL Technologies, cat. no. 07180)-coated plates in mTeSR PLUS (STEMCELL Technologies, cat. no.100-0274) media at 37 °C and 5% CO_2_. Cells were passaged when over 80% confluent. Passaging was performed by washing cells in PBS (Life Technologies, cat. no. 14190-250) and incubating them with ReLeSR (STEMCELL Technologies, cat. no. 05872) for 5 minutes at room temperature, and gently tapping the culture vessels to break hiPSC colonies to smaller colonies for sub-culture.

### Embryoid Body (EB) formation and differentiation to myeloid precursor-like cells

HiPSCs were dissociated to single-cell suspension by incubation with Versene (Life Technologies, cat. no. 15040-033) until cell clumps were not visible. 1×10^4^ cells were plated in mTeSR PLUS media supplemented with 10µM ROCKi (Biogems, cat. no. 1293823), BMP4 (50 ng/ml. R&D Systems, cat. no. 314-BP-010), SCF (20 ng/ml. R&D Systems, cat. no. 255-SC-050), and VEGF (50 ng/ml. R&D Systems, cat. no. 293-VE-010) in ultra-low attachment 384-well U-bottom plates for two days at 37 °C and 5% CO_2_. After two days, 50 µl media without ROCKi were added to each well, and cells were left in culture for two more days. On day 4, EBs were transferred into gelatin-coated T175 flasks (one flask per one 384-well plate) containing 50 ml X-VIVO 15 (Lonza, cat. no. 04-418Q) media supplemented with IL-3 (25 ng/ml. Peprotech, cat. no. 200-03), M-CSF (100 ng/ml. Peprotech, cat. no. 300-25), Glutamax (2mM. Gibco, cat. no. 35050061), and β-mercaptoethanol (55µM. Gibco, cat. no. 21985-023). Four days after EB deposition into the gelatin-coated flasks, a half-media change was carried out. Seven days after EB deposition into flasks (designated as week 0 of myeloid differentiation), a 70 ml-media change was performed. Eleven days after EB transfer into flasks, 40 ml media were added to the culture, and then 110 ml media were collected 14 days after the EB deposition into flasks (designated as week 1 of myeloid differentiation). The 110 ml media collected on week 1 of myeloid differentiation contain myeloid precursor-like cells, however these cells are not used for assays until week 3 of myeloid differentiation, because during those early differentiation timepoints the precursor cells are still immature. The schedule of week 1 is repeated every week until the cultures reach week 10, with 70 ml media added after every precursor-like cell collection and 40 ml media added 4 days after precursor collection, leading to a total volume of 110 ml media collected each week, containing the harvested myeloid precursors.

### HiPSC-derived macrophage differentiation and stimulation

Myeloid precursor-like cells collected from week 3 until week 10 of the differentiation process described above were cultured in RPMI 1640 (Gibco, cat. no. 61870010) media containing 10% FBS (Gibco, cat. no.10099-141) and 100 ng/ml M-CSF for different number of days according to the applications described in this manuscript. For our omics characterisation experiments, cells were cultured in the aforementioned media for 6 days and then stimulated with IFN-γ (20 ng/ml, R&D Systems, cat. no. 285-IF-100) plus LPS (100 ng/ml, MilliporeSigma, cat. no. L5293) or with IL-4 (20 ng/ml, Thermo Fisher, cat. no. PHC0045) for 4h or 24h. Cells were then processed for multi-omics characterisation. For genome editing experiments, macrophages were differentiated for 5 days.

### Culture of primary macrophages

Frozen aliquots of 4 donors of negatively selected CD14^+^ monocytes were purchased from Discovery Life Sciences (DLS). Human biological samples were sourced ethically, and their research use was in accordance with the terms of the informed consents under an IRB/REC approved protocol. Cells were thawed in RPMI 1640 media containing 100 ng/ml MCSF and 10% heat-inactivated FBS. Cells were kept in culture for 6 days and then stimulated with IFN-γ (20 ng/ml) plus LPS (100 ng/ml) or with IL-4 (20 ng/ml) for 4 or 24 hours or left unstimulated before being collected for omics analyses.

### HiPSC-derived and primary macrophage sample preparation for RNA-seq analysis

For both hiPSC-derived and primary macrophages, cells were cultured for 7 days in RPMI 1640 media containing 100 ng/ml MCSF and 10% heat-inactivated FBS. Detachment of hiPSC-derived macrophages was carried out by incubating the cells with 10mg/ml lidocaine solution (MilliporeSigma, cat. no. L5647) in PBS at 37 °C for 5 minutes. Primary macrophages were collected by incubation with TrypLE (Thermo Fisher, cat. no. 12604021) at 37 °C for 15-20 minutes. Both cell types, once collected, were washed with PBS to remove TrypLE or lidocaine solution residue and then resuspended in TRI reagent (Zymo Research, cat. no. R2050-1-200) to produce cell lysates. RNA from cell lysates was isolated using Zymo Direct-zol RNA kits (Zymo Research, cat. no. R2050). Libraries were generated using the Tecan Freedom EVO 200 automated system and Illumina TruSeq Stranded mRNA kit (cat. no. 20020595) following manufacturer recommendations, using starting RNA input of 500ng, fragmentation duration of 6 minutes, and 10 cycles for PCR amplification. Following successful QC, libraries were sequenced using the Illumina NovaSeq 6000 S2 Reagent Kit (101bp paired end) (cat. no. 20028314).

### HiPSC-derived and primary macrophage sample preparation for ATAC-seq analysis

For both hiPSC-derived and primary macrophages, cells were cultured for 7 days in RPMI 1640 media containing 100 ng/ml MCSF and 10% heat-inactivated FBS. Detachment of hiPSC-derived macrophages was carried out by incubating the cells with 10mg/ml lidocaine solution (MilliporeSigma, cat. no. L5647) in PBS at 37 ° C for 5 minutes. Primary macrophages were collected by incubation with TrypLE (Thermo Fisher, cat. no. 12604021) at 37 ° C for 15-20 minutes. For each condition, 100,000 cells were collected and nuclei isolated using ATAC-resuspension buffer (Tris-HCl pH7.4 10 mM, NaCl 3 mM, MgCl_2_ 3 mM, NP-40 0.05%, Digitonin 0.01%, Tween-20 0.1%). Nuclei were then incubated with transposition mixture (Tris-HCl pH7.6 20 mM, MgCl_2_ 10 mM, Dimethyl formamide 20% (MilliporeSigma, cat. no. 227056), transposase 100 mM (Illumina, cat. no. 15027865), 1% digitonin (Promega, cat. no. G9441), 0.1% Tween-20) at 37 ° C for 30 minutes, and finally DNA was isolated and qPCR of transposed fragments was performed. After qPCR amplification, manual assessment of the amplification profiles was carried out, to determine the required number of additional cycles to amplify. See Buenrostro et al. for a detailed explanation of how to properly amplify ATAC-seq libraries [67]. A final qPCR amplification step was performed and DNA was isolated using SPRIselect beads (Beckman Coulter, cat. no. B23318) for DNA size purification. ATAC-seq libraries were sequenced using the Illumina HiSeq High Output v4 kit (75bp paired end).

### HiPSC-derived and primary macrophage sample preparation for proteomics analysis

For both hiPSC-derived and primary macrophages, cells were cultured for 7 days in RPMI 1640 media containing 100 ng/ml MCSF and 10% heat-inactivated FBS. Detachment of hiPSC-derived macrophages was carried out by incubating the cells with 10mg/ml lidocaine solution (MilliporeSigma, cat. no. L5647) in PBS at 37 ° C for 5 minutes. Primary macrophages were collected by incubation with TrypLE (Thermo Fisher, cat. no. 12604021) at 37° C for 15-20 minutes. Cell pellets were then frozen at -80 ° C. Cell pellets were lysed in 2 % SDS buffer for 3 min at 95 °C in a thermomixer, followed by digestion of DNA with benzonase (MilliporeSigma, cat. no. E1014-25KU) at 37°C for 1.5 h. After cell lysis, protein concentration was determined using a BCA assay and samples were further processed through a modified version of the single pot solid-phase sample preparation (SP3) protocol as indicated below.

### MOFA analysis

Pre-processing of single-omics datasets: Proteomics intensities were vsn-normalised using the R/Bioconductor package vsn. Donor and MS-experiment effects were subtracted using the removeBatchEffects function of limma. RNA-seq and ATAC-seq counts were vst-normalised respectively, followed by donor effect subtraction. MOFA integration: MOFA2 was run on the top 1000 most variable features of each omics dataset with the objective to compute 10 latent factors. Enrichment analysis on the weights of the inferred latent factors was performed using the Metabase and MSigDB databases. For constructing the MOFA-informed correlation network, only relationships with a Pearson correlation of at least 0.3 were used.

### Flow cytometry-based QC of macrophage phenotype

Macrophages differentiated for 5 days were collected by washing the cells twice in PBS^-/-Ca2+/Mg2+^ and detachment using TrypLE. Once cells were collected, they were incubated for 10 minutes with Human TruStain FcR (Biolegend, cat. no. 422301) to perform Fc blocking, and then stained for macrophage identity markers for one hour on ice. Cells were analysed on a Cytoflex flow cytometer (Beckman Coulter, cat. no. B53012). Antibodies used to detect macrophage identity marker expression were the following: antibodies against CD45 (Biolegend, cat. no. 304017), CD14 (eBiosciences, cat. no. 17-0149), CD163 (R&D Systems, cat. no. FAB1607G), CD11b (eBiosciences, cat. no. 17-0118-42), CD86 (Biolegend, cat. no. 305411).

### Cytokine secretion assays

Cell culture supernatants collected from IFN-γ+LPS- and IL-4-stimulated macrophages were analysed for cytokine abundance levels. IL-6, CXCL9 and TNF-α release from IFN-γ+LPS-stimulated cells was measured using a cytokine bead array (CBA) ELISA (BD, cat. no. 558276, 558286, 560112). Briefly, supernatants from IFN-γ+LPS-stimulated macrophage cultures were diluted 1:5 in PBS+0.1% BSA and then mixed with a solution of multiplexed CBA capture beads, to achieve around 100 beads per analyte per well. The bead/supernatant mix was incubated at room temperature in the dark on a shaker for 2 hours. After this incubation, the detection antibodies were added with no wash. After a further 2-hour incubation in the dark on a shaker, the assay was read on a Mirrorball laser scanning microplate cytometer (SPT Labtech). CCL13, CCL17, CCL22, IL1Ra and MMP1 release from IL-4-stimulated macrophage cultures was measured using a custom Luminex multiplex cytokine assay (Bio-Techne). Neat (non-diluted) supernatants were assayed in a 384-well plate format using the manufacturer-recommended protocol and read on a Luminex Flexmap 3D instrument.

### Immunofluorescence

After culture supernatant collection, cells were fixed, stained and processed for immunofluorescence imaging.

Briefly, cells were fixed by addition of 4% PFA in PBS for 20 minutes. Cells were then permeabilised using PBS+0.1% Triton-X for 20 minutes, and subsequently blocked using PBST (PBS+0.1% Tween20) supplemented with 2% goat serum for 2 hours.

CD38 (Thermo Fisher, cat. no. MA1-19316) primary antibody solution was added to IFN-γ+LPS-stimulated cells. MAOA (Abcam, Recombinant MAOA antibody [EPR7101], cat. no. ab126751) and TGM2 (Thermo Fisher, monoclonal antibody, CUB 7402, cat. no. MA5-12739) primary antibody solutions were added to IL-4-stimulated cells. Primary antibodies were diluted in blocking solution. Incubations were carried out overnight at 4 ° C.

Following incubation with the primary antibodies, the cells were washed three times for 5 minutes with PBST, and then incubated with fluorophore-conjugated secondary antibodies along with the Hoechst 33258 (Thermo Fisher, cat. no. H21491) and Cell Mask Orange (Invitrogen, cat. no. C10045) dyes. Secondary antibody incubations were carried out for 2 hours at RT in the dark.

Following a final 3 washes with PBST for 5 minutes each, PBS was added to all wells and cells were imaged on an InCell2200 instrument. Images were subsequently analysed on the Columbus software (Revvity) to calculate cell number and marker expression intensities per well.

### Proteomics sample preparation for macrophage protein turnover study

Myeloid precursor-like cells were harvested and cultured in base SILAC macrophage media (Thermo Scientific, cat. no. 88365) supplemented with SILAC-light amino acids (L-Arginine light, MilliporeSigma, cat. no. A8094-25G; L-Lysine light, MIlliporeSigma, cat. no. L9037-25G) for 5 days. To label newly-synthesized proteins, cells were then exposed to SILAC-heavy media for 10 days, and cell samples were harvested every day for analysis. Peptides from pre-existing versus newly-synthesized proteins were distinguished by their mass differences due to incorporation of light or heavy arginine and lysine (L-Arginine heavy, Thermo Scientific, cat. no. 88434; L-Lysine heavy, MilliporeSigma, cat. no. 608041-1G) during downstream mass spectrometric analysis.

For sample harvesting, cells were washed twice in PBS^-/-Ca2+/Mg2+^ and then incubated for 10 minutes in TrypLE at 37 °C. Cells were then lysed in 2% SDS buffer for 3 minutes at 95 °C in a thermomixer, followed by digestion of DNA with benzonase at 37 °C for 1.5 hours.

All samples were processed through a modified version of the single pot solid-phase sample preparation (SP3) protocol as described in ref [68]. Briefly, proteins in 2% SDS were bound to paramagnetic beads (SeraMag Speed beads, GE Healthcare, cat. no. 45152105050250, 651521050502) by addition of ethanol to a final concentration of 50%. Contaminants were removed by washing 4 times with 70% ethanol. Proteins were digested by resuspending in 0.1 mM HEPES (pH 8.5) containing TCEP, chloracetamide, trypsin, and LysC following overnight incubation. Peptides were collected and, after lyophilization, subjected to TMT labelling.

Peptides were labelled with isobaric mass tags (TMT10; Thermo Fisher Scientific, cat. no. 90110) using the 10-plex TMT reagents [69, 70]. The labelling reaction was performed in 100 mM HEPES, pH 8.5 at 22 °C and quenched with glycine. Labelled peptide extracts were combined into a single sample per experiment and lyophilized.

#### Sample preparation for mass spectrometry, prefractionation

Lyophilised samples were resuspended in 1.25% ammonia in water. The whole sample was injected onto a precolumn (2.1 mm × 10 mm, C18, 3.5 μm (Xbridge, Waters)) at a flow rate of 15 μl min−1. Separation was done at 40 μl min−1 on a reversed-phase column (1 mm × 150 mm, C18, 3.5 μm (Xbridge, Waters)) with a 115-min-long gradient ranging from 97% buffer A (1.25% ammonia in water) to 60% B (1.25% ammonia in 70% acetonitrile in water) [71].

#### Liquid chromatography with tandem mass spectrometry

Samples were dried in vacuo and resuspended in 0.05% trifluoroacetic acid in water. Half of the sample was injected into an Ultimate3000 nanoRLSC (Dionex) coupled to a Exploris 480 (Thermo Fisher Scientific). Peptides were separated on custom-made 50 cm × 100 μm (ID) reversed-phase columns (Reprosil) at 55 °C. Gradient elution was performed from 2% acetonitrile to 30% acetonitrile in 0.1% formic acid and 3.5% DMSO over 2 hours. Exploris 480 mass spectrometers were operated with a data-dependent acquisition [72].

#### Peptide and protein identification and quantification

Mascot 2.5 (Matrix Science, Boston, MA) was used for protein identification, in a first search 30 ppm peptide precursor mass and 30 mDa (HCD) mass tolerance for fragment ions was used for recalibration according to Cox et al. [73] followed by search using a 10 ppm mass tolerance for peptide precursors and 20 mDa (HCD) mass tolerance for fragment ions. The search database consisted of a customized version of the SwissProt sequence database (SwissProt Human release December 2018, 42 423 sequences) combined with a decoy version of this database created using scripts supplied by Matrix Science. Carbamidomethylation of cysteine residues was set as fixed modification. Methionine oxidation, and N-terminal acetylation of proteins, and TMT modification of peptide N-termini and Lysine were set as variable modifications.

For SILAC experiments, Carbamidomethylation of cysteine residues was selected as a fixed modification, and the following were selected as variable modifications: oxidation of methionine, acetylation of protein N-termini, SILAC heavy label 13C(6) 15N(4) on arginine and SILAC heavy label 13C(6) 15 N(2) on lysine.

#### Data analysis

Quantification of TMT reporter ions was achieved as described by Savitski et al. [71]. Peptide precursor intensity-based quantification for protein turnover experiments using dynamic SILAC was performed using a modified version of isobarQuant [74] as described in [71, 75]. In short, protein fold changes at different time points were calculated using the intensity ratios of heavy vs. light SILAC peptides and were used for subsequent protein turnover analysis. Protein half-lives were estimated for each protein by fitting a linear model to the time course of the heavy vs. light protein fold changes and determining the time point where this fold change equals 1.

Downstream analyses were performed with R and Bioconductor [76]. TMT reporter intensities were used to calculate the residual KO expression in CRISPR/Cas9-edited hiPSC-derived macrophages over the non-targeting control condition.

### Genome editing in hiPSC-derived macrophages for arrayed CRISPR screen

The arrayed CRISPR screen was performed by RNP delivery into day 5 macrophages using nucleofection. The RNP complex was formed *in vitro* by incubating gRNAs (Synthego) with Alt-R™ S.p. Cas9 Nuclease V3 protein (IDT, cat. no. 1081059) at a 1.2:1 gRNA-to-Cas9 molar ratio at room temperature for 10 minutes. 10μg of Cas9 protein was prepared per reaction, with 250,000 cells.

The gRNA design included commonly 3, occasionally 2 or 1 gRNA(s) targeting the same gene to maximise chances of indel formation and therefore gene editing efficiency. Macrophages were differentiated for 5 days and then lifted as described above, resuspended in P3 buffer (Lonza, cat. no. V5SP-3010) and then mixed with the RNP complex to a final volume of 20μl per reaction. RNP was electroporated into the cells using the 4D Lonza 384-well Nucleofector® System (Lonza, cat. no. AAU-1001) with the DP148 program. Cells were then diluted in macrophage maintenance media and plated into assay plates at a final density of 12,500 cells per 384-well. Plates were left in culture for 6 days to allow genome editing and subsequent protein knock-down effects to take place. Cells were then stimulated for 24 hours before being used in downstream assays. GRNA library is reported in Supplementary Table 1.

### Statistical analysis methods for arrayed CRISPR screen results

The various cytokine and imaging endpoints were analysed in a univariate fashion. Before analysis, data quality control was performed, and it was assessed whether data should be analysed on the original or (log) transformed scale. For most endpoints, analysis on the log transformed scale was recommended.

Next, data was normalised to cell numbers (or log_10_ cell numbers) via the use of robust regression (*ROBUSTREG Procedure in SAS, using Method=MM*) [77]. Using the Stimulated NT control wells, we regressed log_10_ cytokine levels against log_10_ cell counts. Slope and intercept estimates were obtained based on these fits and then applied to data from the entire plate. Separate slope and intercept estimates were obtained for each endpoint and plate.

After normalising the data, hit selection analysis was performed. First, robust mean and robust standard deviations were calculated for the Stimulated and Unstimulated NT controls. This was again done separately for each endpoint and plate. The approach used to obtain robust estimates is described in [78]. Once the robust estimates were obtained, different hit selection criteria and cutoffs as well as Z-Prime quality control measures were calculated based on these robust estimates. In addition, we also calculated the Strictly Standardized Mean Difference (SSMD) metric [33]. SSMD aims to provide better control of false positive and false negative rates, and it is calculated as follows:

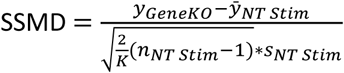

where y_GeneKO_ is the transformed and normalised cytokine or imaging endpoint value of a specific GeneKO well, *y̅_NT Stim_* and s_NT Stim_ are the robust mean and robust standard deviation estimates of the Stimulated NT control wells on each plate, K = n_NT Stim_ – 2.48 and n_NT Stim_ is the sample size of the Stimulated NT control wells on each plate.

Hit identification was primarily based on SSMD values, and a cutoff of -/+2 was used for downregulation or upregulation, respectively.

### Genome editing in hiPSC-derived macrophages for pooled CRISPR screen

The protocol used to perform genome editing for the pooled screen is a modified version of the protocol reported above for the arrayed screen. Myeloid precursor-like cells were differentiated to macrophages for 5 days. To deliver the gRNA library and achieve integration of a single lentiviral copy into the cells, cells were transduced with lentivirus aiming at 30% transduction efficiency and incubated overnight, following media change on day 6. The virus dilution factor was determined in advance, by performing transductions with serial dilutions of the lentivirus solution and assessing transduction efficiencies by flow cytometry 5 days post-seeding. Cas9 protein was electroporated into the cells on day 7, in P3 buffer, on the 4D Lonza electroporator unit, using programme DP148. In this case, 60 pmol of Cas9 protein was electroporated per 250,000 cells. To enrich the infected cell population, flow cytometry sorting based on ZsGreen expression was carried out on day 12. On day 14 of macrophage differentiation, cells were stimulated using IFN-γ (20 ng/ml) and LPS (100 ng/ml). After 24 hours, cells were detached and processed for single-cell RNA-seq with gRNA capture analysis using the 10x Genomics Chromium Platform.

In parallel, wild type cells from the same hiPSC donors were cultured for 14 days and then stimulated with IFN-γ (20 ng/ml) and LPS (100 ng/ml), or left unstimulated for 24 hours, and then collected for single-cell RNA-seq analysis.

The seeding densities of the gene-edited and wild type control cells used after electroporation were reduced to avoid over-confluency of the cultures. 40,625 cells/cm^2^ or 36,458 cells/cm^2^ were used for the electroporated or wild type control cells, respectively.

### CROP-seq gRNA design

GRNAs were designed using CRISPick [39]. For each gene KO, 4 gRNAs were designed and included in the CROP-seq library. 8 gRNAs were designed as non-targeting/non-cutting controls and 8 gRNAs were designed as cutting controls within the AAVS1 locus.

All target DNA sequences are reported in Supplementary Table 2.

### CROP-seq gRNA cloning and lentivirus generation

GRNA oligos were synthesized and cloned into the modified CROP-seq vector described by Catalinas et al. [35] by Vector Builder. Lentivirus carrying the CROP-seq-gRNA vectors was also sourced by Vector Builder. The pooled gRNA library was constructed from 3.8 x 10^5^ single colonies, providing over 100-fold coverage of the intended gRNAs. Paired-end sequencing with a read length of 150 bp was performed on an Illumina NovaSeq instrument. Read 1 data was aligned with a reference gRNA list, which successfully identified 80 out of the 80 designed gRNAs - providing 100% coverage.

### CROP-seq in hiPSC-derived macrophages

Single-cell RNA-seq libraries were generated using a Chromium NextGEM Single Cell 3’ HT Reagent Kit (10x Genomics, cat. no. CG000416) according to the manufacturer’s protocol. Taking advantage of donor pooling, we overloaded (cell input 60,000 / lane) the Chromium chip to recover 35,000 cells per lane. Whole-transcriptome libraries, prepared from captured cells, were indexed using Dual Index Plate TT Set A (10X Genomics).

To enhance the detection of sgRNA(s) expressed in a given cell, sgRNA amplicons were enriched from 10x cDNA following a method adapted from Hill *et al.* [79]. Briefly, using 10ng of non-fragmented 10x cDNA, PCR was performed with primers described in Table 1 using GoTaq PCR Master Mix (Promega, cat. no. A6001). To avoid over-amplification, we performed parallel PCR reactions with varying number of cycles and retained the reaction that provided optimal quantity of amplicon with least number of cycles (12 cycles). Using one-tenth of the purified PCR product, we performed the second nested PCR round (9 cycles). Again, using one-tenth of the purified round 2 PCR product, a sequencing-ready sgRNA amplicon library was prepared by 6 cycles of PCR using primers from Dual Index Plate TN Set A (10x Genomics).

**Table 1.**
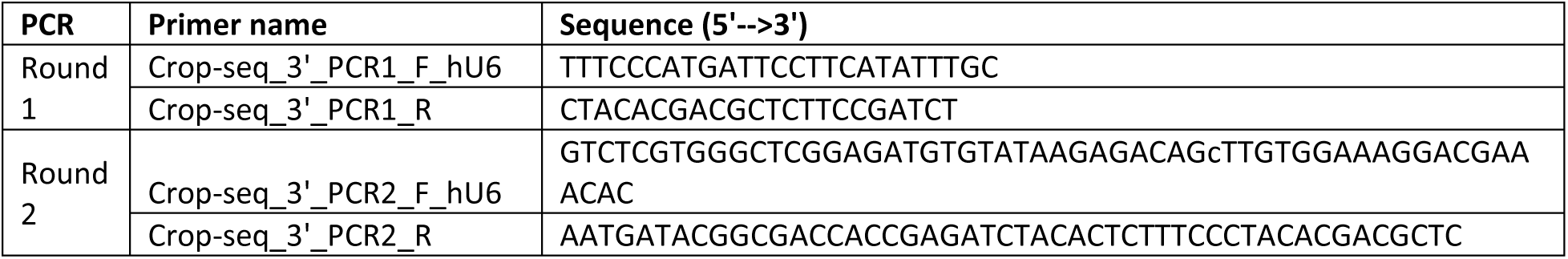

Single-cell whole-transcriptome and sgRNA amplicon libraries were mixed at a 7:1 molar ratio and sequenced for 28:10:10:90 cycles (read1:index1:index2:read2) using a NovaSeq 6000 system (Illumina).

### CROP-seq data processing

Single-cell sequencing data were processed using cellranger count v6.0 with the --include-introns parameter. The transcriptome was mapped against the human reference genome provided by 10X Genomics (hg38, Ensembl annotation version 98; https://cf.10xgenomics.com/supp/cell-exp/refdata-gex-GRCh38-2020-A.tar.gz). A feature reference file containing all gRNA sequences present in the library was supplied to generate counts for each gRNA in the gRNA library. We processed 10 technical replicates, each of which represents a different channel of a 10x chip. Each replicate contains cells from each of the three donors and together covers both transduction and stimulation states. We recovered a median of 20,000 cells from each technical replicate with 38,912 mean reads and 4,452 mean genes per cell. We observed a slight increase in UMI counts recovered from Unstimulated vs. Stimulated conditions (median 41 vs. 28; Supplementary Figure 3). Guide RNA calling was performed using a custom script to assess which cells were transduced with which gRNA. Expected gRNA proportions were calculated from sequencing of the gRNA library and were provided to aid the calling process. The filtered_feature_matrix produced by cellranger were imported into R for downstream analysis.

#### Donor demultiplexing

Donor demultiplexing was performed on all samples using Souporcell [80] to assess which cells were derived from the same hiPSC donor. Souporcell was also used to compare the pooled donor samples from this experiment to previously-generated donor-specific samples to label each donor with its appropriate name.

#### Quality control

Analysis was conducted using bioconductor Single Cell Experiment (SCE) architecture in R. Cells were removed if they were called as a doublet according to either DonorDemux or DoubletDetection (30% of cells), or if they could not be assigned to any donor (4%) - this reduced the number of cells from 197,086 to 133,932 cells. We noticed that doublets from DoubletDetection had large library sizes, whereas doublets identified through SNP-based DonorDemux had smaller library sizes. The 133,932 cells were then filtered down to 118,239 by removing cells with high MT expression, or low gene detection/total counts assessed through scater [81] by using the isOutlier function on a per sample basis setting nmads=3 and an overall minimum gene detection threshold of 2,500. 13,050 of these cells were derived from the “WT” control samples, whereas 105,189 cells were derived from the gRNA-transduced samples.

#### gRNA assignment

To ascertain the successful transduction of each gRNA in each cell, a binomial test was employed on UMI counts derived from the gRNA library. The binomial test considered the cell’s total library size, accounting for the anticipated proportions of each gRNA, to establish the minimum UMI count necessary to classify a gRNA as present above background noise. The anticipated proportions of each gRNA were estimated through DNA sequencing of the lentiviral library used for transduction. Any gRNAs exhibiting a Bonferroni-adjusted p-value less than 0.001 were deemed present in the respective cell. Furthermore, significant gRNA assignments supported by three or fewer UMI counts were excluded. In total, gRNA calls were confidently assigned to 77,345 cells (74%) including 49,842 cells (47%) with a single gRNA assignment, which were used as the basis for further analysis. In total, we obtained a median of 110 cells per gRNA per donor for the unstimulated samples, and 66 cells per gRNA per donor for the stimulated samples (Supplementary Figure 3).

## Key resource table

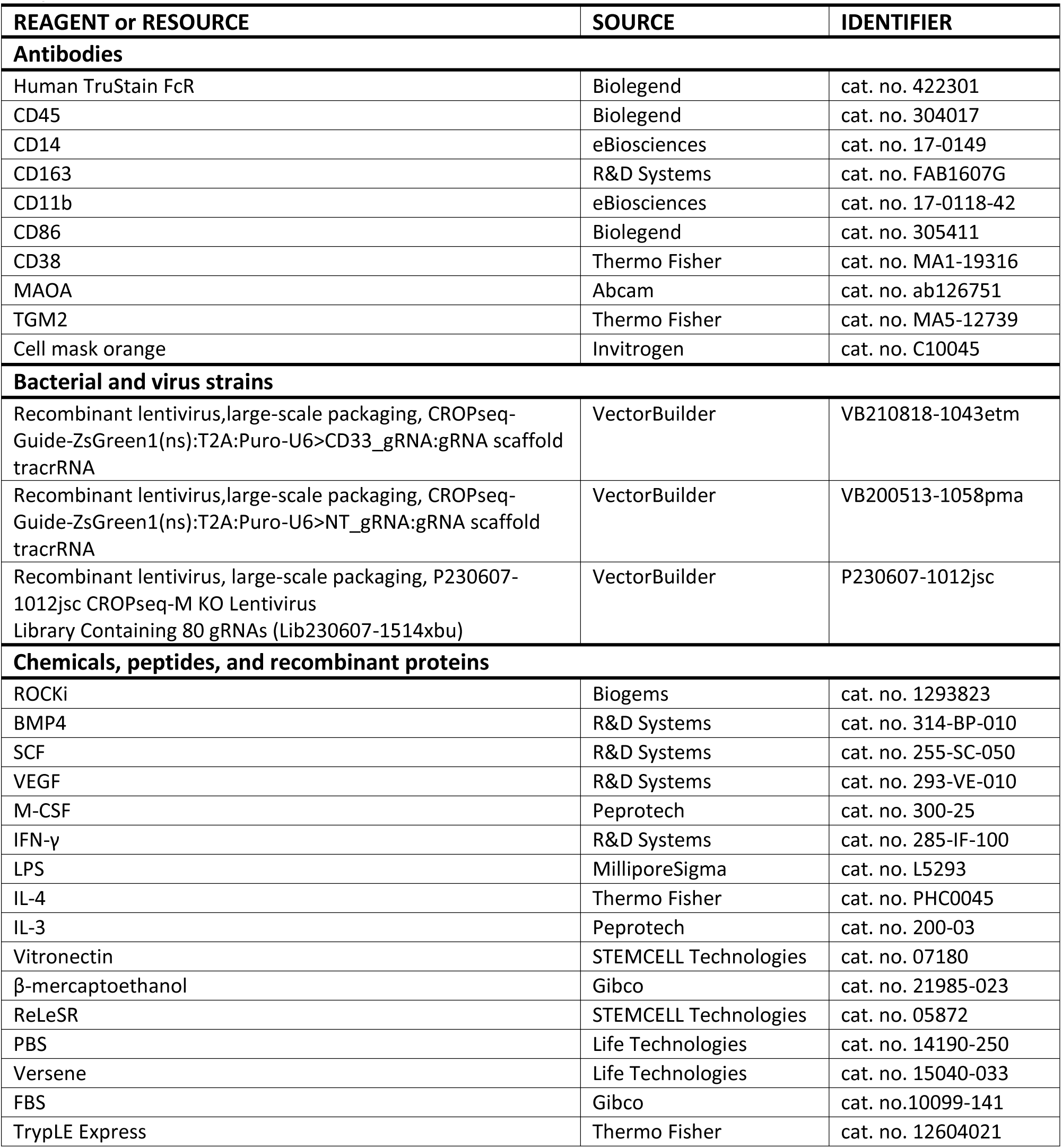
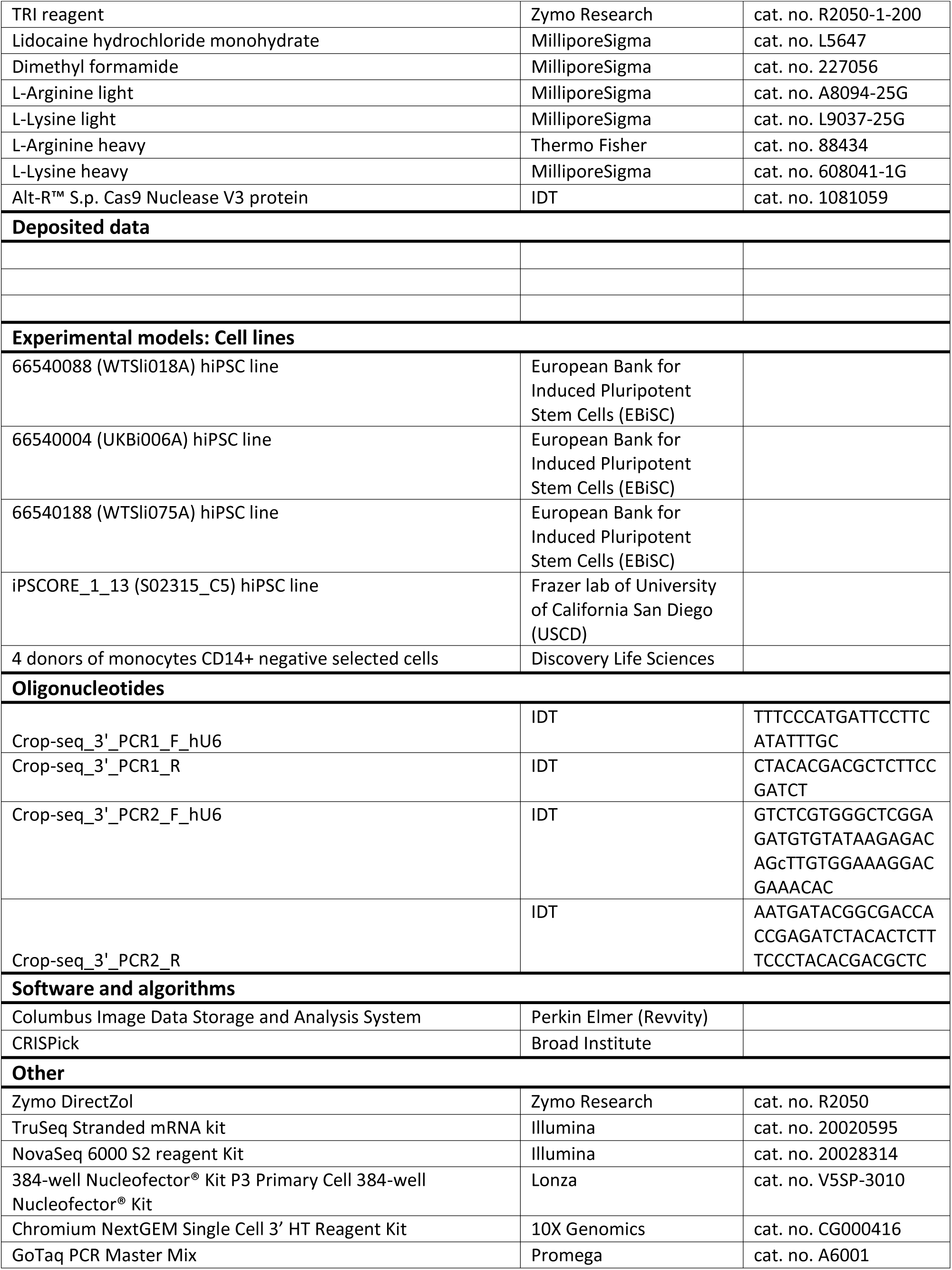
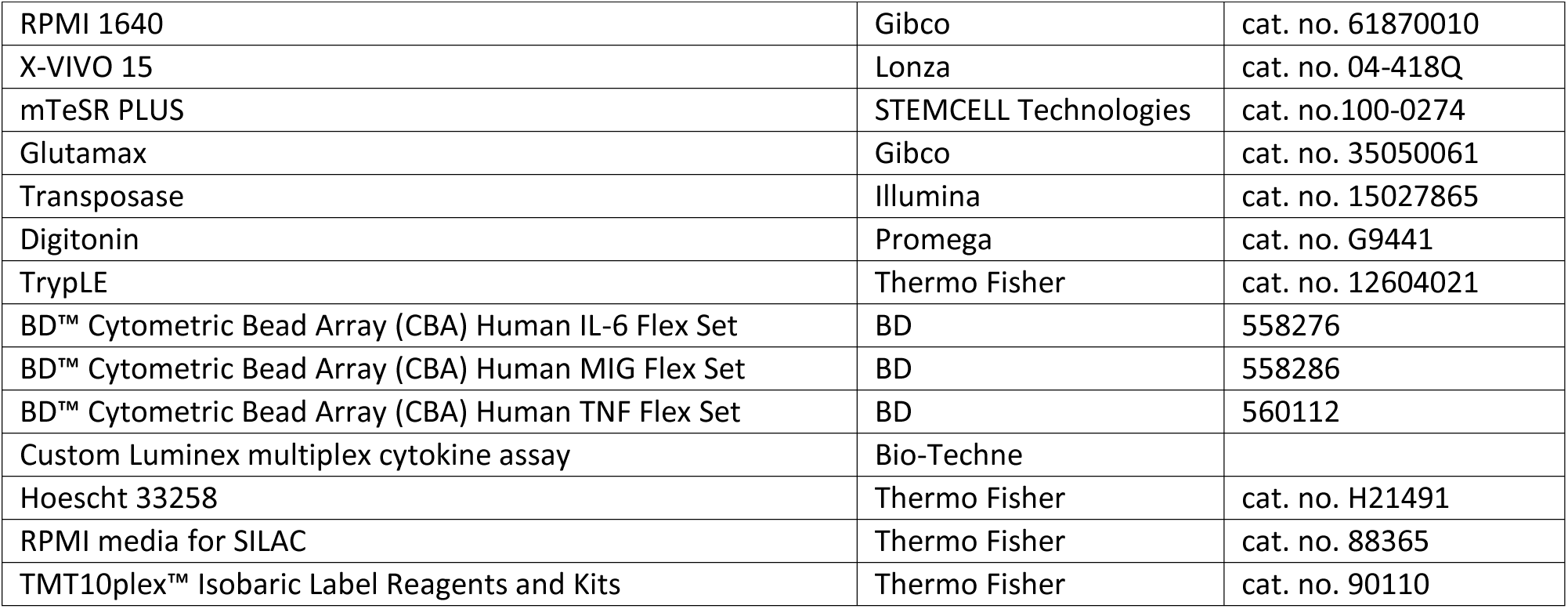

## Competing Interest Statement

All authors are employees of GSK, a global healthcare company that may conceivably benefit financially through this publication.

## Supplementary Figures

**Supplementary Figure 1.**
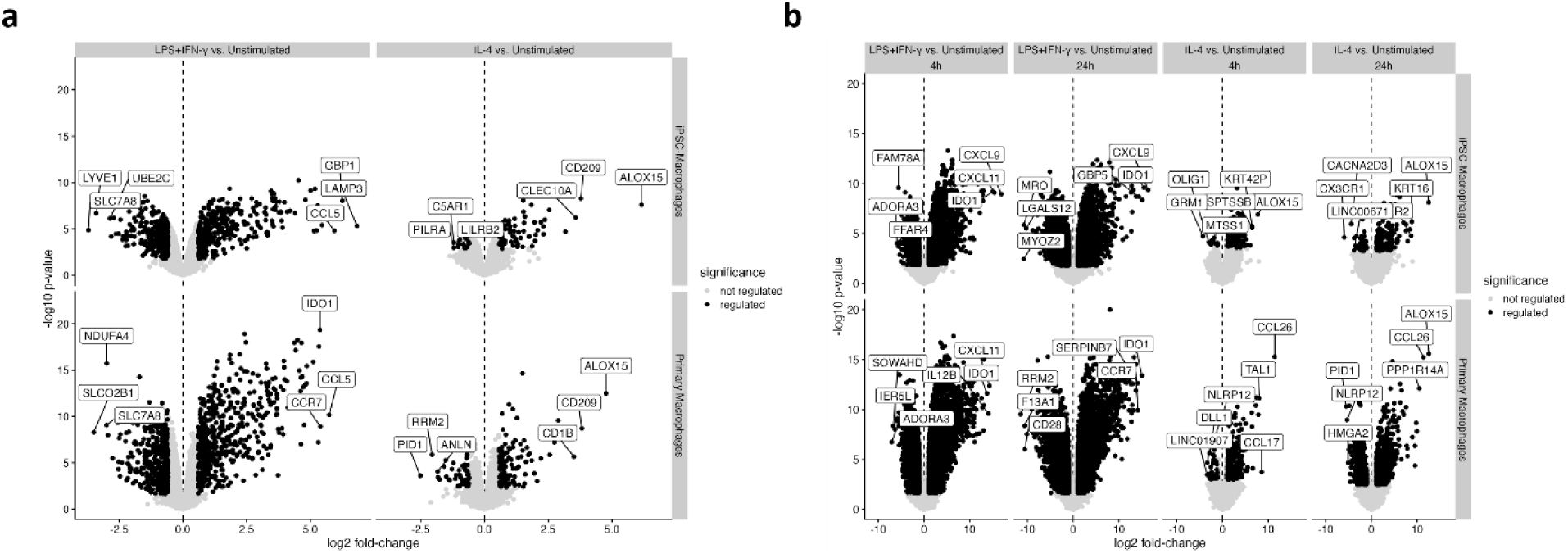
Differential expression analysis results for proteomics and transcriptomics data on hiPSC-derived and primary macrophages after stimulation with IFN-γ+LPS or IL-4. a) Volcano plots showing the differential expression analysis results for proteomics data on hiPSC-derived macrophages derived from 3 donors and primary macrophages derived from 4 donors, either left unstimulated or treated with IFN-γ (20 ng/ml) + LPS (100 ng/ml) or with IL-4 (20 ng/ml). Samples for proteomics analysis were collected 24h after stimulation in both cases. Top 3 upregulated and downregulated proteins are highlighted in the plots for each condition. b) Volcano plots showing the differential expression analysis results for RNA-seq data on hiPSC-derived macrophages derived from 3 donors and primary macrophages derived from 4 donors, either left unstimulated or treated with IFN-γ (20 ng/ml) + LPS (100 ng/ml) or with IL-4 (20 ng/ml). Samples for transcriptomics analysis were collected 4h and 24h post-stimulation in both cases. Top 3 upregulated and downregulated transcripts are highlighted in the plots for each condition.

**Supplementary Figure 2.**
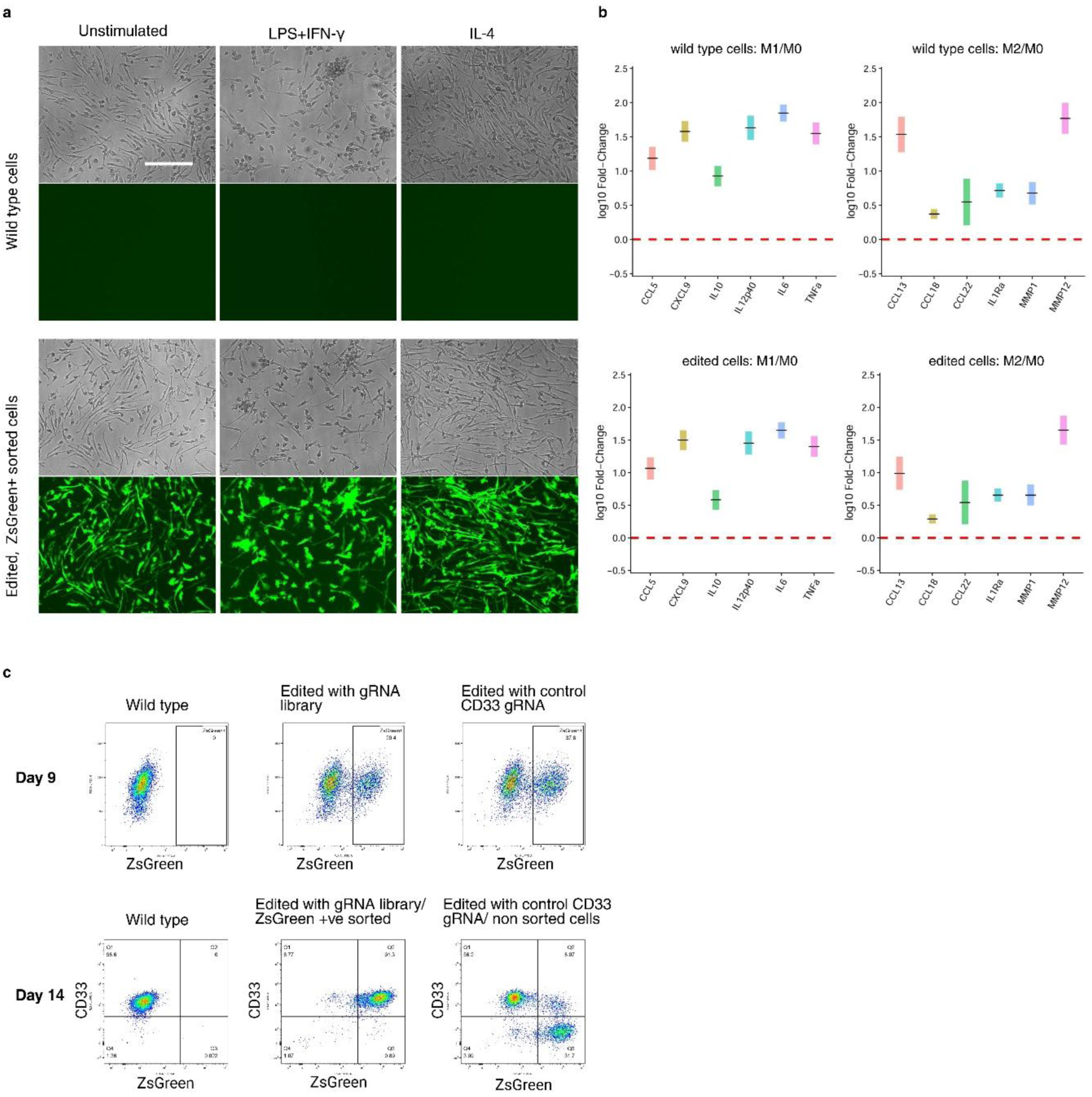
Characterisation of the functional phenotype of macrophages genome-edited using the SLICE protocol. a) HiPSC-derived macrophages on day 5 of differentiation from myeloid precursor-like cells were transduced with lentiviruses carrying our modified CROP-seq vector, and subsequently sorted for ZsGreen expression on day 12 of the differentiation protocol. Stimulated and unstimulated cells were imaged on day 15 to assess ZsGreen expression as a proxy for gRNA expression. Image scale bars represent 300μm in length. b) Wild type (top plots) or lentivirus-transduced (bottom plots) macrophages were stimulated with IFN-γ+LPS (left plots) or with IL-4 (right plots) for 24h, and cell culture supernatants were subsequently analysed for Type 1 or Type 2 inflammatory cytokine release, respectively. c) Macrophages that were processed using the SLICE workflow were analysed by flow cytometry at two different timepoints. Day 9 flow cytometry analysis plots show ZsGreen expression in macrophages transduced with a vector containing CD33 gRNAs or with the full gRNA vector library. Day 14 flow cytometry analysis plots show ZsGreen expression (X axis) against CD33 protein expression (Y axis), with the latter being used as a control demonstrating successful genome editing achieved by our CROP-seq method at the protein level.

**Supplementary Figure 3.**
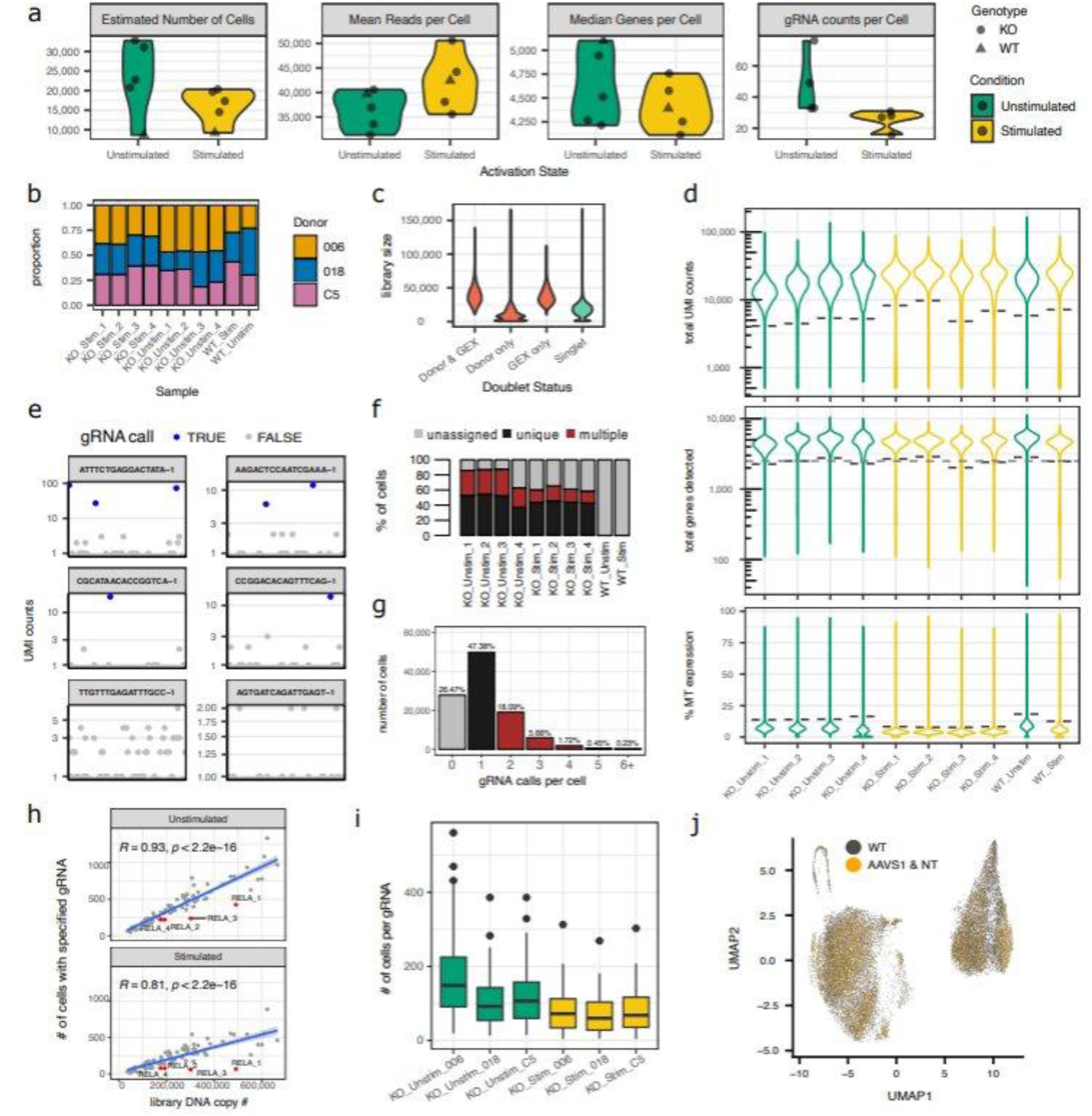
Processing of CROP-seq data. A) Cell and read recovery. Key sequencing QC metrics for CROP-seq technical replicates (channels of a 10x Genomics chip) coloured by activation state (Unstimulated, green; Stimulated, yellow; colour scheme used throughout). Samples with no gRNA transduction are denoted by triangles. Unstimulated samples show a modest increase in recovered UMI counts relative to Stimulated samples. B) Donor deconvolution. Bar plot showing the proportion of cells assigned to each hiPSC donor for each CROP-seq technical replicate. C) Doublet removal. Donor-based demultiplexing identifies doublets through the presence of RNA derived from multiple donors, while gene-expression (GEX)-based methods assess whether a cell’s transcriptional profile resembles a combination of two randomly selected cells. These two doublet classes are largely non-overlapping and show differing library size distributions. Cells identified as doublets (red) were removed, while singlets (blue) were retained. D) Removal of poor-quality cells. Distributions of total UMIs per cell, number of detected genes per cell, and fraction of reads mapping to mitochondrial genes, used for cell-level QC filtering. Cells were removed if they were 3 median absolute deviations (MADs) greater than median mitochondrial (MT) expression, or 3 MADs lower than the median gene detection or total counts, or if they did not meet a minimum gene detection threshold of 2,500, indicated by dashed lines. E) Guide RNA calling. GRNA expression for randomly selected cells with either multiple gRNAs detected per cell (top), a single unique gRNA detected (middle), or no gRNA detected (bottom). F) Proportions of gRNA call categories across technical replicates: unique (black), multiple (red), and unassigned (grey). G) Distribution of gRNA calls per cell. Cells with only a single unique gRNA call (47%) were used as the basis for downstream analysis. H) Scatter plot showing the relationship between the relative abundance of each gRNA in the plasmid library assessed by DNA sequencing (X axis) versus the number of cells uniquely called for that gRNA assessed by scRNA-seq (Y axis). GRNAs targeting RELA are highlighted given their under-representation in the scRNA-seq data compared with expectations based on their relative abundance in the DNA library. I) Bar plot showing the number of cells assigned per gRNA for each donor. Median recovery was 110 cells per gRNA per donor under Unstimulated conditions and 66 cells per gRNA per donor under Stimulated conditions. J) UMAP showing the localisation of cells carrying control gRNAs (AAVS1 and non-targeting [NT]) compared with unedited wild type (WT) cells.

## Author contributions

M.M. experimental lead, writing and conceptualisation; E.K. experimental lead (CROP-seq method) and writing; T.P. data analysis and figure generation; T.A. arrayed screen data generation; A.G., B.C. data generation, figure generation, writing; K.R., A.Q.M., W.P., D.V., S.G., H.F., F.Z. data analysis; C.H. data generation; N.Z. data generation and analysis; M.A., D.T. data generation; B.S., M.M. experimental lab support; A.D.T., L.E., N.S., M.L.; L.M. writing and conceptualisation.

## Acknowledgements

We would like to thank Alejandro Armesilla Diaz, Sara Schmidt, Linda Myers, Jamie Ifkovits, Amy Yeung for their support in cell generation; David Brierley and Kirsty Ford for their contribution in the arrayed screen; David Mayhew, George Royal, Carl Fishwick, Wendy Halsey, Melchert Stephanie, Traini Christopher, Van Horn Stephanie, Stephanie Lehman, Johanna Korben, Mathias Kalxdorf for their support in omics data generation, Megan Ermler for her immunology expertise. Project has been funded solely by GSK.

## Declaration of generative AI and AI-assisted technologies in the writing process

During the preparation of this manuscript the authors used Jules, an internal GSK tool, in order to improve readability of the manuscript. After using this AI tool, the authors reviewed and edited the content as needed, and take full responsibility for the content of the published article.

## Resource availability Lead contacts

Requests for further information and resources should be directed to and will be fulfilled by the lead contacts, Matteo Martufi and Lisa Mohamet.

## Material availability

Materials used in this manuscript are captured in the key resource table and in the method section. This study did not generate unique reagents.

## Data availability

- Bulk RNA-seq data presented in Figure 2 have been deposited at GEO at GEO: **GSE341821** and are publicly available as of the date of publication.
- ATAC-seq data presented in Figure 2 have been deposited at GEO at GEO: **GSE342643** and are publicly available as of the date of publication.
- Single-cell RNA-seq data presented in Figures 5 and 6 have been deposited at GEO at GEO: **GSE341348** and are publicly available as of the date of publication.

